# Mitochondrial volume fraction and translation speed impact mRNA localization and production of nuclear-encoded mitochondrial proteins

**DOI:** 10.1101/529289

**Authors:** Tatsuhisa Tsuboi, Matheus P. Viana, Fan Xu, Jingwen Yu, Raghav Chanchani, Ximena G. Arceo, Evelina Tutucci, Joonhyuk Choi, Yang S. Chen, Robert H. Singer, Susanne M. Rafelski, Brian M. Zid

## Abstract

Mitochondria are dynamic in their size and morphology yet must also precisely control their protein composition according to cellular energy demand. Although nuclear-encoded mRNAs can be localized to the mitochondrial outer membrane, the importance of this localization in altering mitochondrial protein composition is unclear. We have found that, as yeast switch from fermentative to respiratory metabolism, there is an increase in the fraction of the cytoplasm that is mitochondrial. This drives the localization of certain nuclear-encoded mitochondrial mRNAs to the surface of the mitochondria. Through tethering experiments, we show that mitochondrial mRNA localization is necessary and sufficient to increase protein production to levels required during respiratory growth. Furthermore, we find that ribosome stalling impacts mRNA sensitivity to mitochondrial volume fraction and counterintuitively leads to enhanced protein synthesis by increasing mRNA localization to the mitochondria. This points to a mechanism by which cells are able to use translation elongation and the geometric constraints of the cell to fine-tune organelle-specific gene expression through mRNA localization while potentially circumventing the need to directly coordinate with the nuclear genome.

## INTRODUCTION

Mitochondria are essential cellular organelles that are key sources of ATP generation via oxidative phosphorylation as well as the assembly of iron-sulfur clusters and many other catabolic and anabolic reactions (Attardi and Schatz, 1988). To support mitochondrial function, thousands of nuclear-encoded proteins are imported into mitochondria from the cytoplasm (Morgenstern et al., 2017). This has to be coordinated with the gene expression of the mitochondrial genome, which in *Saccharomyces cerevisiae* contains 13 genes (Borst and Grivell, 1978). While cells can generate ATP through mitochondrial oxidative phosphorylation, they can also use glycolysis as an alternative means of generating ATP. *Saccharomyces cerevisiae*, for example, switches its metabolism from respiration to glycolysis/fermentation in response to oxygen depletion or increasing fermentable sugar levels (De Deken, 1966). This metabolic change is known to dramatically change the mitochondrial morphology (Egner et al., 2002). The protein content of yeast mitochondria also shows dynamic changes in response to shifting cellular energy demands (Morgenstern et al., 2017; Paulo et al., 2016). Oxidative phosphorylation protein coding mRNAs are known to gradually increase their protein synthesis as the growth environment changes from fermentative growth to respiratory conditions (Couvillion et al., 2016).

mRNA localization is a means to post-transcriptionally regulate gene expression at both a temporal and spatial level (Martin and Ephrussi, 2009). In the 1970s, electron microscopy analysis found that cytoplasmic ribosomes can be localized along the mitochondrial outer membrane (Kellems et al., 1974). Recent microarray and RNA-seq analyses of biochemically fractionated mitochondrial membranes and fluorescent microscopy analysis have identified subsets of nuclear-encoded mRNAs that are mitochondrially localized (Fazal et al., 2019; Gadir et al., 2011; Garcia et al., 2007; Marc et al., 2002; Saint-Georges et al., 2008; Williams et al., 2014). It has been shown that both the 3′ UTR and coding regions, primarily through mitochondrial targeting sequences (MTSs), contribute to mitochondrial localization. One class of localized mRNAs (Class I) was shown to be dependent on the Puf3 RNA-binging protein through binding motifs in the 3′ UTR (Saint-Georges et al., 2008). Another class of mRNAs was localized to the mitochondria independently of Puf3 (Class II). Many of the localized mRNAs show reduced association upon polysome dissociation through EDTA or puromycin treatment, implicating translation as a necessary factor for mRNA localization (Eliyahu et al., 2010; Fazal et al., 2019). The mitochondrial translocase of the outer membrane (TOM) complex has been shown to impact mRNA localization through interaction with the nascent MTS (Eliyahu et al., 2010), while the outer-membrane protein OM14 has been shown to be a mitochondrial receptor for the ribosome nascent-chain-associated complex (NAC) (Lesnik et al., 2014). Isolating mitochondrially localized ribosomes to perform proximity-specific ribosome profiling revealed that many mitochondrial inner-membrane protein mRNAs are co-translationally targeted to the mitochondria (Williams et al., 2014). These observations have suggested a mechanism of co-translational protein import into mitochondria for a subset of nuclear-encoded mitochondrial mRNAs.

While mRNA localization is a way to control gene expression and there is strong evidence for the localization of mRNAs to the mitochondria, the use of this mRNA localization to alter the composition of mitochondria in different environmental conditions has not been explored. Furthermore, we know that mitochondria are very dynamic structurally in relation to the metabolic needs of the cell, yet how this changing morphology may directly impact mRNA localization and gene expression has also not been investigated. Here we report that interactions between mitochondria and mRNA/nascent-peptide (MTS) complexes can be altered by both the kinetics of protein synthesis and the fraction of cytoplasm that is mitochondrial, leading to condition-dependent mitochondrial mRNA localization during respiratory conditions. This localization subsequently leads to enhanced translation and protein expression for these condition-specific localized mRNAs during respiratory conditions.

## RESULTS

### mRNA association to mitochondria differs between fermentative and respiratory conditions

To explore how mRNA localization is impacted by the metabolic state of the cell and changing mitochondrial morphologies, we developed a methodology to quantify mRNA location and mitochondrial 3D structure in living cells. To do this, we visualized mitochondria using the matrix marker Su9-mCherry and single-molecule mRNAs with the MS2-MCP system every 3 seconds in a microfluidic device (Figures 1A and S1). We reconstructed and analyzed the spatial relationship between the mRNAs and mitochondria using the ImageJ plugin Trackmate (Tinevez et al., 2017) and MitoGraph V2.0, which we previously developed to reconstruct 3D mitochondria based on matrix marker fluorescent protein intensity (Rafelski et al., 2012; Viana et al., 2015) (Figure 1A). We measured the distance between mRNA and mitochondria by finding the closest meshed surface area of the mitochondrial matrix (Figure S2, STAR Methods).

**Figure 1.**
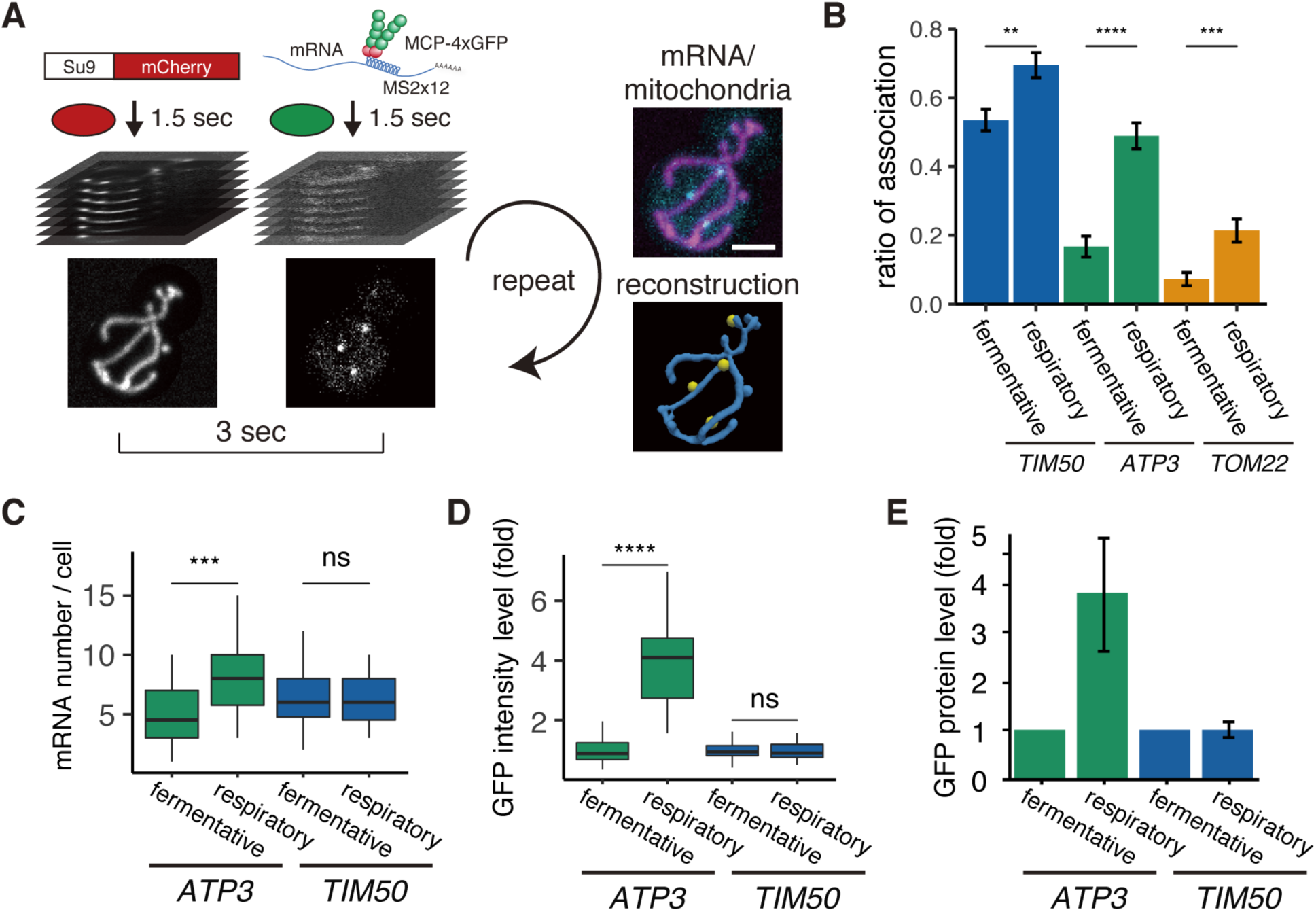
mRNA Association to Mitochondria Differs between Fermentative and Respiratory Conditions. (A) Experimental setup for live imaging. Mitochondria were visualized by Su9-mCherry and mRNAs were visualized by the single molecule MS2-MCP tethering system. Z-stacks were taken within 1.5 sec for each individual channel, and multiple Z-stacks were merged into a series. (Right top) Z-projected image of a live cell. Cyan: mRNA, purple: matrix. Scale bar, 2 μm. (Right bottom) Reconstructed mRNA and mitochondria. Yellow: mRNA, blue: mitochondria. (B) The ratio of the mitochondrially associated mRNA per cell (n>27) of the different mRNA species in fermentative and respiratory conditions. Error bar represents standard error of the mean (s.e.m.). (C) *ATP3* mRNA and *TIM50* mRNA expression number per cell. MCP-GFP foci were counted per cell (n>27). (D) Atp3p-GFP and Tim50p-GFP fusion protein expression level using GFP fluorescent intensity per cell. (E) Atp3p-GFP and Tim50p-GFP fusion protein expression level using western blotting with anti-GFP antibody. Error indicates standard deviation of three independent experiments. (C–E) Statistical significance was assessed by Mann–Whitney U-test (**** P < 0.0001; *** P < 0.001; ** P < 0.01; ns, no significant difference).

We first analyzed three different mRNAs (Saint-Georges et al., 2008; Williams et al., 2014). Two mRNAs have previously been found to be mitochondrially localized and contain a mitochondrial targeting sequence (MTS): *ATP3* mRNA, which encodes the gamma subunit of ATP synthase, and *TIM50*, which encodes a component of the inner membrane translocase. The third mRNA, *TOM22*, encodes an outer-membrane translocase that does not contain an MTS and has previously been found to be predominantly diffusely localized (Gadir et al., 2011; Garcia et al., 2010; Williams et al., 2014). During fermentative growth, we observed *TIM50* mRNA to be strongly associated with the mitochondria, whereas *TOM22* showed low association consistent with previous studies (Figures 1B and S2, and STAR Methods).

Even though *ATP3* had previously been categorized as a mitochondrially localized mRNA (Gadir et al., 2011; Saint-Georges et al., 2008), we unexpectedly found this to be condition-dependent as it has low association with mitochondria, similar to *TOM22*, in fermentative conditions. However, during respiratory conditions it strongly shifted towards association with the mitochondrial surface, in a manner more similar to *TIM50* (Figure 1B). This means that nuclear-encoded mitochondrial mRNAs do not have to be solely mitochondrially localized or diffusely localized; instead, they can show a switch-like behavior depending on the metabolic needs of the cell.

There are large changes in mitochondrial composition as yeast switch from fermentative to respiratory metabolism (Morgenstern et al., 2017). To investigate whether there may be a relationship between mRNA localization and gene expression, we measured mRNA levels via number of mRNAs per single cell and protein levels via both single-cell measurements and bulk assays in both fermentative and respiratory conditions. For the condition-dependent *ATP3* mRNA, we found that protein levels increased 4-fold, whereas mRNA levels increased less than 2-fold when cells were grown in respiratory versus fermentative conditions. *TIM50* mRNA, which is constitutively localized to the mitochondria under both conditions, showed no change in protein or mRNA levels in respiratory conditions (Figures 1C–1E). This suggests a relationship between mRNA localization to the mitochondria and protein production.

### Relationship of mRNA localization to mitochondrial volume fraction

As yeast cells shift to respiratory conditions, the mitochondrial volume increases while the cell cytoplasmic volume decreases, thus leading to an increase in the mitochondrial volume fraction in respiratory conditions (Egner et al., 2002) (Figures 2A and S3). While *ATP3* mRNA showed a strong condition-dependent localization, *TIM50* and *TOM22* mRNAs also showed modestly increased mitochondrial association during respiratory conditions (Figure 1B). We wondered what impact the reduction in the availability of free cytoplasmic space due to mitochondrial expansion had on mRNA co-localization, especially for *TOM22*, which is not known to bind to the mitochondria. To test this, we quantified both the mitochondrial localization of each mRNA and changes in mitochondrial volume fraction at a single-cell level. We found that *TOM22* showed a linear increase in co-localization that was directly proportional to mitochondrial volume fraction (Figure 2B). We also found that *ATP3* mRNA was more sensitive to mitochondrial volume fraction than *TIM50* and *TOM22*. This sensitivity was independent of nutrients, as fermentative yeast cells showed a larger increase in *ATP3* localization as the mitochondrial volume fraction increased (Figure 2B). At the lowest mitochondrial volume fractions, *ATP3* localization was similar to the diffusely localized *TOM22* mRNA, whereas at the highest volume fractions, its localization was close to the mitochondrially localized mRNA *TIM50*. This suggests that increased mitochondrial volume fraction drives *ATP3* mRNA localization to mitochondria.

**Figure 2.**
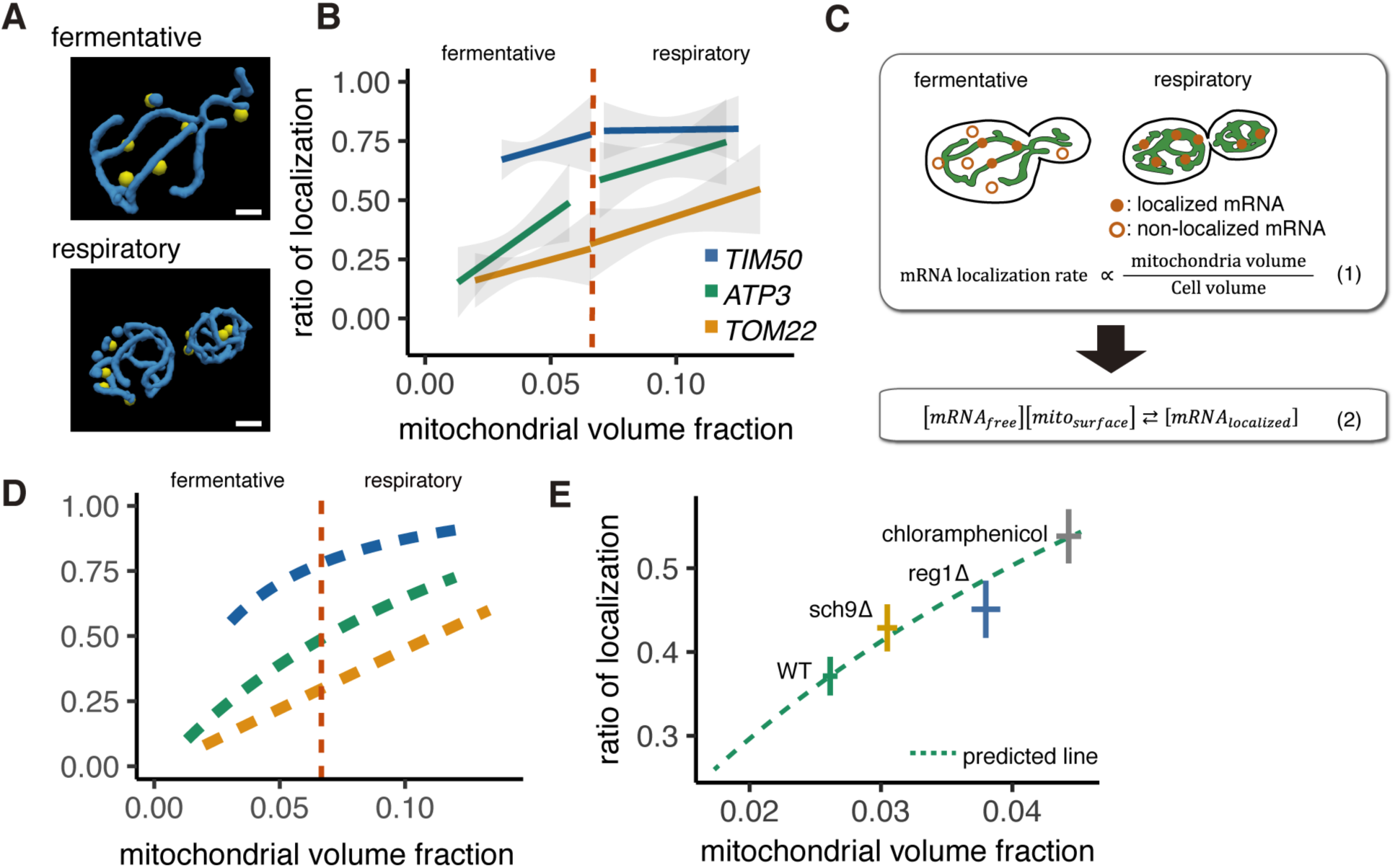
Mitochondrial Volume Fraction Correlates with mRNA Localization. (A) Snapshot of reconstructed mitochondrial surface (blue) and *TIM50* mRNA foci (yellow) in fermentative and respiratory conditions. Scale bar, 1 μm. (B) Relationship between the mitochondrial volume fraction and the ratio of mRNA localization to mitochondria. Trend line was depicted according to the best linear fit of the ratio of localization and mitochondrial volume fraction of single cells in each condition of different mRNAs (n>27). Dotted red line marks the difference between fermentative and respiratory conditions for mitochondria volume fraction. Gray region surrounding the trend lines represents the 95% confidence interval (CI) for each line. (C) Schematic of *in silico* experiment. (Top) Brownian particle distribution indicates that mRNA localization rate correlates with mitochondrial volume fraction. (Bottom) Hypothetical thermodynamic equilibrium of binding of mRNA to mitochondria. [mRNA_localized_], mitochondrial localized mRNA; [mRNA_free_], free diffusing mRNA; [mito_surface_], mitochondrial surface where mRNA can bind. (D) Mitochondrial volume fraction and mRNA localization have stoichiometric correlation. Relationship of ratio of mRNA localization to mitochondria and mitochondrial volume fraction from mathematical modeling was plotted. Yellow line represents linearly fitted line for *TOM22* mRNA. Green and blue lines were plotted through equilibrium constant of 2.4K_0_ and 8.8K_0_, respectively, as described in Methods. (E) In glucose conditions, *sch9*Δ and *reg1*Δ mutant strains as well as chloramphenicol addition exhibit higher mitochondrial volume fraction and increase the localization of *ATP3* mRNA to the mitochondria (n>27). Predicted relationship of ratio of mRNA localization to mitochondria and mitochondrial volume fraction from mathematical modeling of WT strains was plotted as a green dotted line. Mean values of each axis for WT, mutant cells, and cells with chloramphenicol addition were plotted as crosses. Error bar represents s.e.m.

To further test our hypothesis that mRNA localization is regulated by mitochondrial volume fraction, we designed *in silico* experiments based on our experimentally measured cell and mitochondrial boundaries and used a mathematical model to investigate how particles of varying affinities would co-localize with mitochondria (Figure 2C). We were able to recapitulate the behavior of *TOM22* via a model of an ideal Brownian particle with no affinity for the mitochondria, which showed linearly correlated localization with mitochondrial volume fraction (Figure 2C, equation (1); Figures 2D and S4A). We then set up a simple equilibrium equation where the baseline equilibrium constant, K_o_, was set by a freely diffusing particle like *TOM22* and multiplied by the affinity, A, of the particle for the mitochondria, thus giving K = AK_o_ (Figure 2C, equation (2); STAR Methods). As the value of A increased in the simulation, the ratio of mitochondrial localization of the mRNA for a given mitochondrial volume fraction increased as well (Figure S4B). We then applied this relationship to estimate the experimental values of A to be 2.4 and 8.8 for *ATP3* and *TIM50*, respectively (Figures 2D and S4C). Notably, mitochondrial volume fraction correlates with mRNA localization, but cell volume and mitochondrial volume alone did not show any correlation through fermentative and respiratory conditions (Figures S4C–S4E). Interestingly, this simple mathematical relationship also recapitulates the shape of the curves in Figure 2B, suggesting that the apparent shift in association that occurs during the switch from fermentative to respiratory conditions may be solely a result of the combination of the mitochondrial volume fraction and the strength of mRNA-specific association and, therefore, not due to the difference in growth condition or to other mechanisms.

To further test this model, we sought to manipulate mitochondrial volume fraction in a multitude of ways. For example, Reg1 is a protein necessary for glucose repression in yeast, and *reg1*Δ cells exhibit increased mitochondrial function in glucose media (Adachi et al., 2017; Hubscher et al., 2016). Supporting our model, we observed increased mitochondrial volume fraction in rich glucose conditions in *reg1*Δ mutant cells as well as increased *ATP3* mRNA localization to the mitochondria (Figures 2E, S5A–S5D, and S6). Additionally, the antibiotic chloramphenicol is known to decrease mitochondrial protein synthesis while having no effect on cytoplasmic translation and has been found to increase the fraction of mitochondrial protein in *Neurospora crassa* and mitochondrial volume in *Tetrahymena pyriformis* (Brody, 1992; Gleason et al., 1975). We sought to test whether chloramphenicol would alter the mitochondria volume fraction in yeast during rich glucose conditions when mitochondrial respiration is not required. We observed an ∼70% increase in mitochondria volume fraction in yeast cells treated with chloramphenicol and, simultaneously, a ∼50% increase in *ATP3* mRNA mitochondrial association during vegetative conditions (Figures 2E, S5A–S5D, and S6). These results support the hypothesis that mitochondrial volume fraction is the important variable for *ATP3* mRNA localization and not a factor related to respiration, as chloramphenicol decreases respiratory function (Williamson et al., 1971) while increasing mitochondrial volume fraction and *ATP3* localization.

Finally, as mitochondrial volume fraction can be increased by either increasing mitochondrial volume or decreasing cytoplasmic volume, we tested whether decreasing cytoplasmic volume could increase *ATP3* mRNA mitochondrial association. *sch9Δ* was previously found to be one of the smallest strains from a genome-wide screen for cell size (Jorgensen et al., 2002). We found that *ATP3* mRNAs showed increased mitochondrial association in *sch9Δ*, though from initial analysis this was not accompanied by an increase in mitochondrial volume fraction. While imaging these cells, we noticed a large increase in the relative vacuole in *sch9Δ* cells; the vacuole volume fraction was ∼6x higher in *sch9Δ* cells versus WT cells. As the vacuole restricts the accessible cytoplasmic volume, we recalculated the mitochondrial volume fraction relative to the accessible cytosol and found that *sch9Δ* cells have a significant increase in mitochondrial volume fraction after this correction (Figures S5E, S5F, and S6). With this corrected mitochondrial volume fraction, we found that, similar to *reg1*Δ and chloramphenicol-treated cells, *sch9Δ* cells had increased localization of *ATP3* mRNA to the mitochondria that was not significantly different either from our experimental measurements relating mitochondrial volume fraction and mRNA association in WT cells or from what would be predicted from our mRNA localization simulation based solely on changing mitochondrial volume fraction (Figure 2E, S5G–S5I, and S6). These results show that the relationship between mitochondrial volume fraction and mRNA localization holds across highly varied perturbations, including nutritional (glucose/glycerol), genetic (*reg1*Δ, *sch9*Δ), and pharmacological (chloramphenicol). These data point to mRNA association with mitochondria that can be tuneable to permit a switch-like transition in mitochondrial localization, as is seen for *ATP3* mRNA (Figure 1B), due to a nutrient-induced change in mitochondrial volume fraction.

### Translation regulates mRNA localization

Given these results, we wanted to delve further into the mechanism of this varying localization and protein production. Even though the mitochondrial protein import machinery is well described, an ER-like signal recognition particle dependent mechanism of co-translational protein import has not been identified for the mitochondria (Golani-Armon and Arava, 2016; Reid and Nicchitta, 2015). However, a series of biochemistry and microscopy analysis showed that some nuclear-encoded mitochondrial protein mRNAs are translated on the mitochondrial surface (Gadir et al., 2011; Garcia et al., 2007; Gold et al., 2017; Kellems et al., 1974; Marc et al., 2002; Saint-Georges et al., 2008; Williams et al., 2014). We therefore investigated the effects of the MTS and of protein translation on mRNA association to mitochondria. We replaced the MTS of Tim50p with an ER-localization signal or introduced an ER-targeting signal at the N-terminus of Tom22p (Wu et al., 2016). Even though *TIM50* mRNA was associated with mitochondria, *ER-TIM50, TOM22*, and *ER-TOM22* mRNAs were not associated with mitochondria (Figure 3A), indicating that the *TIM50* MTS is necessary to recruit this mRNA to mitochondria. To further support the role of the MTS in mRNA localization, we tested whether reducing ribosome-nascent chain association by using the translation initiation inhibitor lactimidomycin (LTM) would affect mRNA localization. We found that *TIM50* mRNA in all conditions and *ATP3* mRNA in respiratory conditions decreased localization to the mitochondrial surface upon LTM addition, while *ATP3* mRNA in fermentative conditions showed only minimal changes in localization upon LTM addition (Figure 3A). These results suggest that the actively translating ribosome drives mRNA localization to mitochondria through production of the nascent N-terminal MTS.

**Figure 3.**
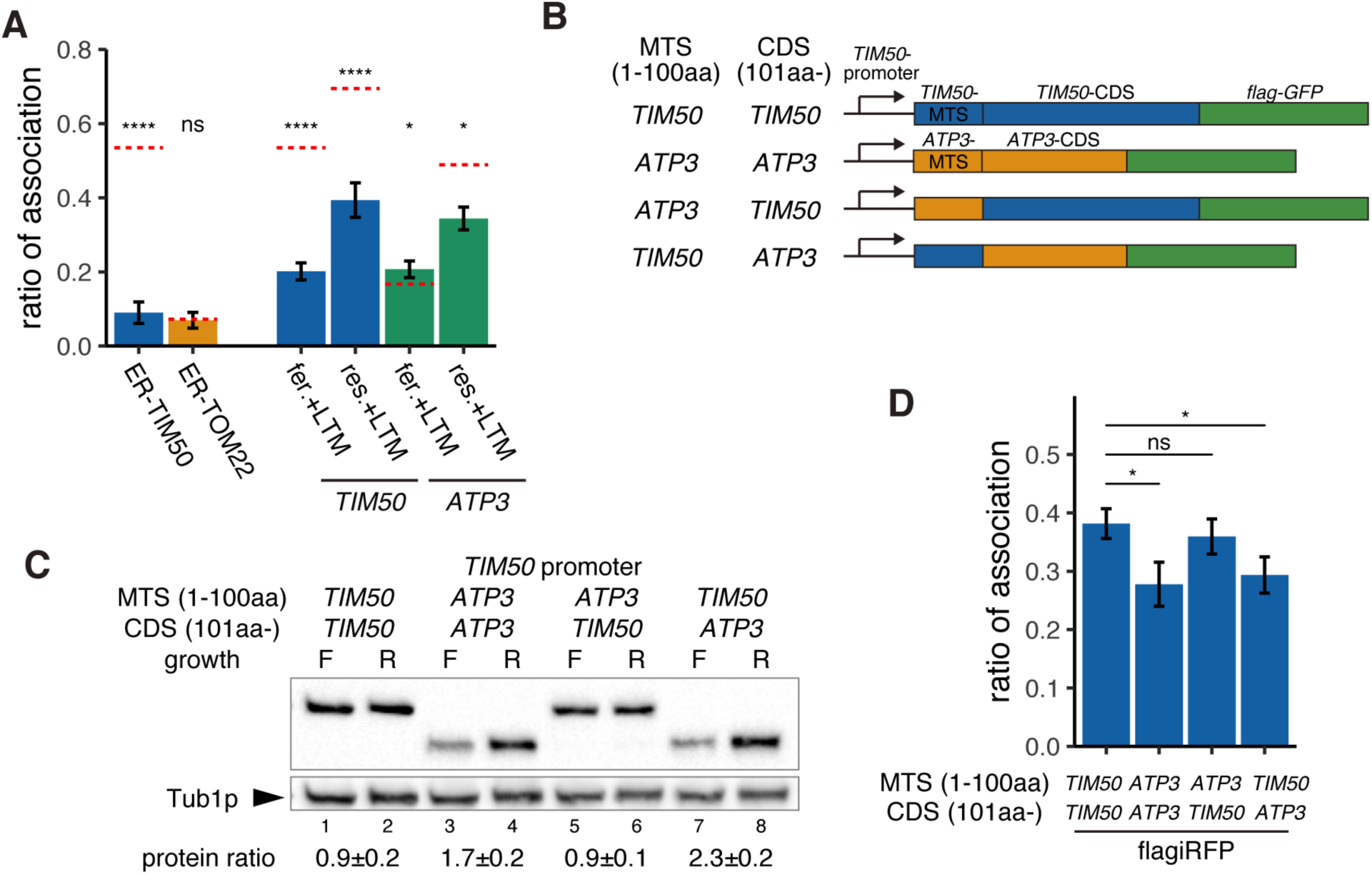
Increased Protein Synthesis and mRNA Localization Is Regulated by the Downstream Coding Sequence. (A) An ER-localization signal and translation inhibitor drugs alter the ratio of the mitochondrial associated mRNA per cell of the strains in Figure 1B (n>20). LTM, 50μM for 20 min. Error bar represents s.e.m. Statistical significance compared with control (value in Figure 1B, red dotted line) was assessed by Mann–Whitney U-test (**** P < 0.0001; * P < 0.05; ns, no significant difference). (B) Schematic of chimeric reporter genes for swapping of MTS (1-100aa) and CDS (101aa-) between *TIM50* and *ATP3*. (C) Protein expression from reporter genes depicted in (B). Growth ‘F’ represents fermentative and ‘R’ represents respiratory conditions. Tub1p was used as internal loading control. Protein expression ratio between respiratory and fermentative conditions is shown in the bottom row. Error indicates standard deviation of three independent experiments. (D) The ratio of mitochondrial associated mRNA per cell of reporter mRNAs in fermentative and respiratory conditions (n>34). Error bar represents s.e.m. Statistical significance was assessed by Mann–Whitney U-test (* P < 0.05; ns, no significant difference).

From our simulation, we were able to recapitulate our experimental observation that *TIM50* mRNA has higher affinity for the mitochondria than *ATP3* mRNA, making its localization less dependent on mitochondrial volume fraction. As the MTS was necessary for localization to the mitochondria, our initial hypothesis was that Tim50p MTS has a higher affinity for the mitochondria than the Atp3p MTS, causing the differences in mRNA affinities. To test this hypothesis, we designed chimeric reporter genes wherein we swapped the MTS sequences between Tim50p and Atp3p under *TIM50* promoter control (Figure 3B). Surprisingly, we found that the downstream coding sequence (CDS) was what differentiated *TIM50* from *ATP3*, not the MTS. When the reporter gene contained the TIM50-CDS, it showed uniform protein production in fermentative versus respiratory conditions, independent of which MTS was present (Figure 3C, lane 1 vs. 2 and lane 5 vs. 6). However, when the reporter gene contained the ATP3-CDS, it showed decreased protein production in fermentative conditions (Figure 3C, lane 3 vs. 4 and lane 7 vs. 8). Similarly, the reporter genes that harbored the ATP3-CDS also showed decreased mitochondrial mRNA association ratios in fermentative conditions (Figure 3D). These experiments suggest that the *TIM50* and *ATP3* MTS have similar affinities for the mitochondria and, importantly, what drives the condition-specific differences in mRNA localization and protein production between these mRNAs is encoded in the downstream CDS.

### Ribosome stalling is important for constitutive mitochondrial localization

Our model proposes that the reason *ATP3* mRNA increases localization in respiratory conditions is that the increased mitochondrial volume fraction increases the probability that the nascent MTS will interact with the mitochondrial surface. If the *ATP3* and *TIM50* MTS have similar affinity for the mitochondria, we hypothesized that the reason *TIM50* has higher mitochondrial association at lower mitochondrial volume fractions is because the downstream CDS increases the chance of association between mitochondria and MTS, possibly by slowed translation elongation. Upon further examination, we found that the *TIM50* CDS has 7 consecutive polyprolines approximately 60 amino acids downstream of the MTS. Polyproline stretches have been shown to mediate ribosome stalling, and, when we investigated a ribosome profiling data set, we found that ribosomes accumulate at this polyproline stretch during fermentative conditions (Zid and O’Shea, 2014) (Figure S7). This suggests a possible mechanism, similar to what has been seen for SRP recognition, by which local slowdown of ribosomes increases the chance that the mitochondria will recognize the *TIM50* MTS and consequently promote its association with the mitochondrial surface (Chartron et al., 2016). To test this, we deleted these polyproline residues and found this caused *TIM50* to be more sensitive to environmental conditions as it reduced both protein synthesis and mRNA localization of *TIM50* during fermentative conditions (Figures 4A–4C). In contrast, the ATP3 coding sequence does not have any obvious strong ribosome stalling sequence. This suggests that *ATP3* mRNA localization and protein synthesis are regulated solely in a manner dependent on mitochondrial volume fraction. If this is true, artificially slowing the ribosomes in fermentative conditions should drive *ATP3* mRNA to become mitochondrially localized. To test this hypothesis, we inserted 7 consecutive polyprolines at 100 amino acids downstream of the start codon and analyzed protein expression in fermentative conditions (Figure 4A, right). We observed that protein production is increased 1.6-fold (Figure 4D) and mRNA localization is increased as well (Figure 4E). These results further suggest that ribosome stalling leads to mRNA localization to mitochondria and increased protein production.

**Figure 4.**
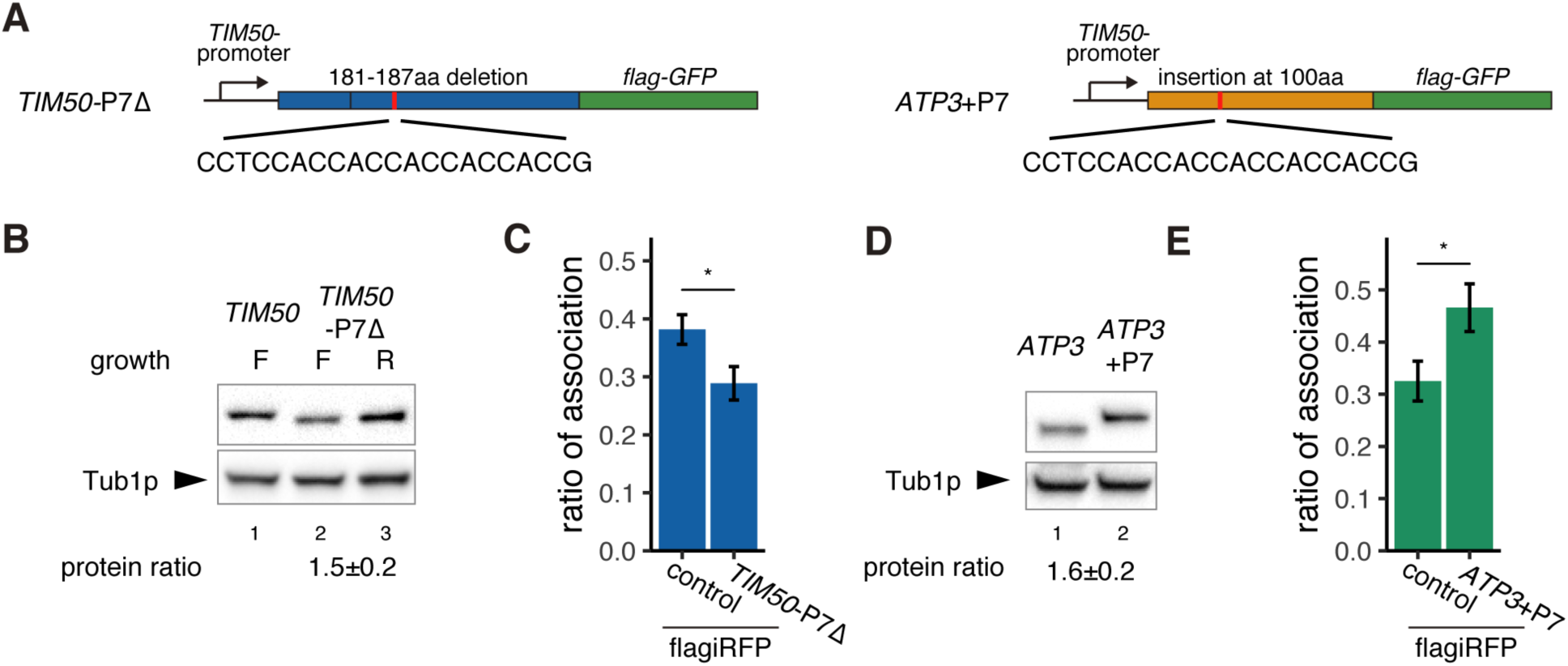
Decreased Translational Elongation Localizes mRNA to Mitochondria. (A) Schematic of deletion of polyproline sequence from *TIM50-GFP* reporter gene and insertion of polyproline sequence into *ATP3-GFP* reporter gene. This construct is called *TIM50*-P7Δ and *ATP3*+P7. (B) Protein expression from reporter genes *TIM50* and *TIM50*-P7Δ. Growth ‘F’ and ‘R’ correspond to fermentative and respiratory conditions, respectively. Tub1p was used as internal loading control. Protein expression ratio between respiratory and fermentative conditions is shown in the bottom row. Error indicates standard deviation of three independent experiments. (C) The ratio of the mitochondrial associated mRNA per cell (n>20) of the reporter mRNAs in fermentative and respiratory conditions. (D) Protein expression from reporter genes *ATP3* and *ATP3*+P7 in respiratory condition. Tub1p was used as internal loading control. Protein expression ratio between the reporter genes is shown in the bottom row. Error indicates standard deviation of three independent experiments. (E) The ratio of the mitochondrial associated mRNA per cell (n>29) of the reporter mRNAs in fermentative conditions.

To further explore whether slowing translation elongation stabilizes the mRNA-ribosome complex with the MTS, thereby giving it more time to associate with mitochondria, we measured mitochondrial mRNA localization following the addition of the translation elongation inhibitor cycloheximide (CHX) (Figure 5A). As our hypothesis predicted, we observed a 3-fold increase in the association of *ATP3* mRNA with mitochondria during fermentative conditions but no increase in the case of *TOM22* mRNA (Figure 5A). Interestingly, CHX treatment only slightly increased *TIM50* mRNA localization. This potentially suggests that a portion of these mRNAs are incompetent for binding, perhaps due to nuclear localization or being in the midst of degradation, and that the majority of competent *TIM50* mRNAs were already localized to the mitochondria.

**Figure 5.**
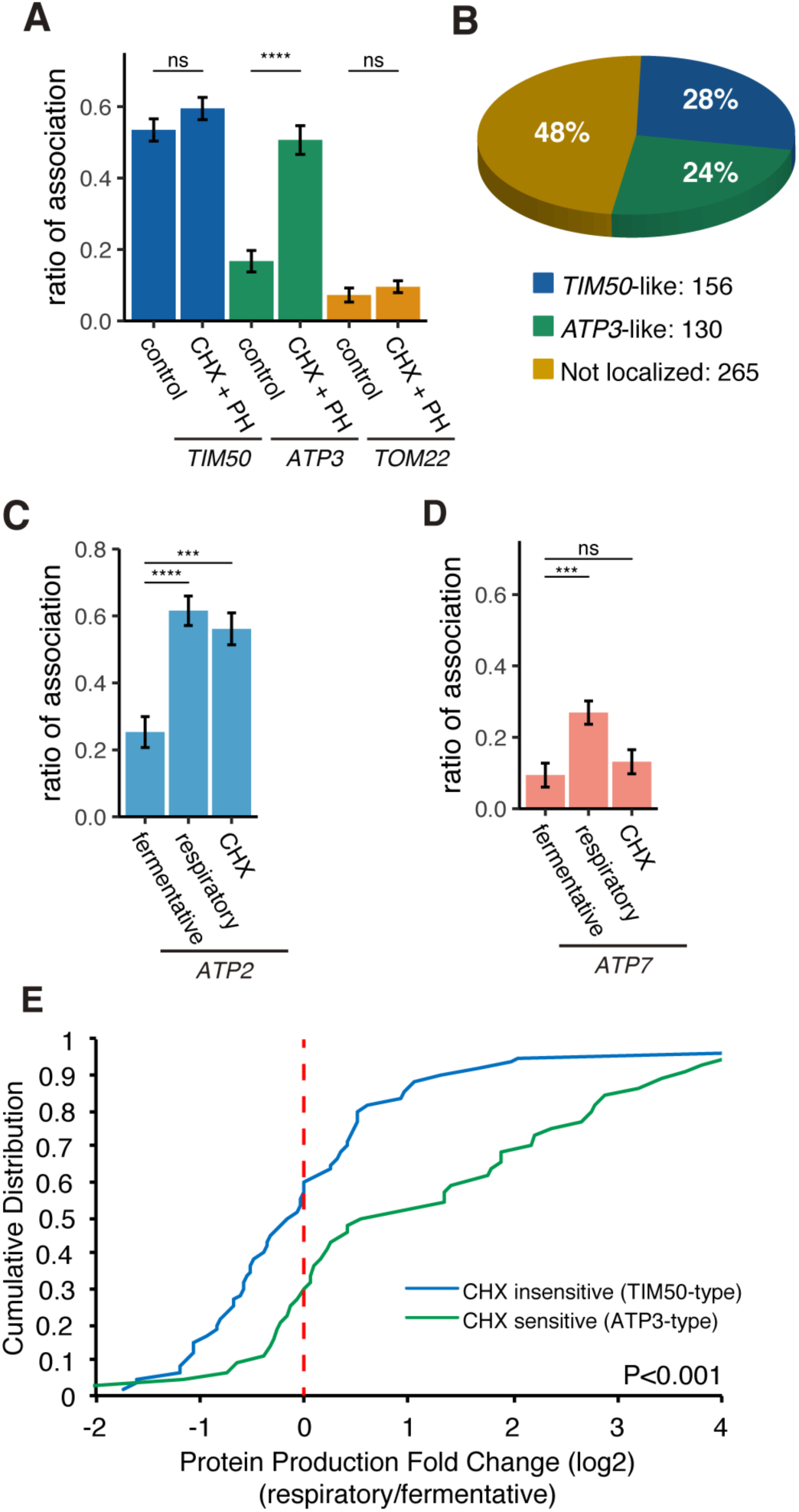
Decreased Translational Elongation Localizes mRNA to Mitochondria. (A) Translational inhibitor drugs alter the ratio of the mitochondrial associated mRNA per cell (n>43) of the strains in Figure 1C. CHX+PH indicates 100μg/mL cycloheximide and 200μg/mL 1,10-Phenanthroline for 10min. Error bar represents s.e.m. (B) Percent distribution of *TIM50* and *ATP3* mRNA mitochondrial localization behavior in annotated mitochondria protein encoding genes. We re-summarized and classified *TIM50*-type mRNA as enriched at the mitochondria regardless of CHX addition and *ATP3*-type mRNA as enriched at the mitochondria only upon CHX addition using proximity ribosome profiling data of annotated mitochondria protein encoding genes (Mitop2) (Elstner et al., 2009; Williams et al., 2014). All other mRNAs were considered not localized. (C, D) The ratio of the mitochondrial associated mRNA per cell (n>16) of the different mRNA species in fermentative, respiratory, and CHX-treated conditions. Error bar represents s.e.m. Statistical significance in this figure was assessed by Mann–Whitney U-test (**** P < 0.0001; *** P < 0.001; ** P < 0.01; * P < 0.05; ns, no significant difference). (E) Protein production is increased upon a switch from fermentative to respiratory conditions in ATP3-type genes. Cumulative distribution of protein production (Couvillion et al., 2016) of TIM50-type (n=60) and ATP3-type genes (n=44) were depicted with blue and green lines, respectively. Statistical significance in this figure was assessed by Student’s t-test.

A previous study found that 130 of 551 annotated nuclear-encoded mitochondrial mRNAs are sensitive to translation elongation rate and become localized to the mitochondrial surface upon cycloheximide treatment, similar to *ATP3* (Figure 5B) (Williams et al., 2014). Interestingly, all of the ATP synthase subunits that are conserved from bacteria to eukaryotes are sensitive to cycloheximide, except for the ε subunit, *ATP16*, whereas all of the nonconserved subunits are insensitive to cycloheximide (Table S3). We wondered whether this sensitivity may be indicative of mRNAs that also switch their localization as the mitochondrial volume fraction increases during respiratory conditions (Table S3). We tested *ATP2*, the conserved β subunit of the ATP synthase, and found that it was similar to *ATP3* in that it showed a large increase in localization according to volume fraction change upon a shift to respiratory conditions and was sensitive to cycloheximide (Figures 5C and S8). However, the nonconserved subunit, *ATP7*, behaved more like *TOM22* in that it was insensitive to cycloheximide and had a much smaller increase in mitochondrial localization in respiratory conditions than *ATP2* or *ATP3* (Figure 5D). We then wanted to more globally explore the connection between sensitivity to translation elongation and changes in gene expression during the metabolic shift from fermentation to respiration when mitochondrial volume fraction dramatically increases. We focused on Class II mRNAs that were found to be localized to the mitochondria during respiratory conditions independently of Puf3 (Saint-Georges et al., 2008) and subdivided these into *ATP3*-type mRNAs that were cycloheximide-sensitive in localization to mitochondria during fermentative conditions and *TIM50*-type that were constitutively localized to mitochondria in fermentative conditions (Williams et al., 2014). We used previously generated ribosome profiling data on yeast cells under glucose and glycerol conditions (Couvillion et al., 2016). From this data we found that *ATP3*-type mRNAs had a significant increase in their protein productive capacity versus *TIM50*-type as they had >2-fold more ribosomes engaged in translation in the shift from glucose to glycerol (Figure 5E). The increase in protein production was caused by both a significant increase in translation efficiency and mRNA levels for *ATP3*-type mRNAs during respiratory conditions (Figure S9). From an independent proteomics data set under glucose and glycerol conditions, we also found that *ATP3*-type genes had an increase in protein abundance during respiratory conditions versus *TIM50*-type genes (Figure S9) (Morgenstern et al., 2017). These results point to translational elongation sensitivity and condition-dependent localization being general strategies to fine-tune gene expression of certain nuclear-encoded mitochondrial genes.

### mRNA localization to the mitochondria drives enhanced protein synthesis

Recently, it has been shown that there is active cytoplasmic translation on the mitochondrial surface in *Drosophila* and that a subset of nuclear-encoded mitochondrial proteins are translationally regulated by the localization of specific RNA-binding proteins to the mitochondrial surface (Zhang et al., 2016). We therefore hypothesized that mRNA localization to mitochondria may be a way to drive the coordinated increase in mitochondrial protein production observed in respiratory conditions. An alternative explanation is that increased translation drives more nascent protein production, which increases mRNA localization to the mitochondria. To directly test these two possibilities, we analyzed the effect of driving mRNA localization to mitochondria on protein expression. To accomplish this, we tethered reporter mRNAs to mitochondria by MS2 sequences. We inserted the MCP protein into the C-terminus of Tom20p and Tom70p, two well-characterized proteins on the outer mitochondrial membrane, and analyzed subsequent protein production (Figure 6A). We found that tethering *TIM50-flag-GFP* and *ATP3-flag-GFP* mRNA to the mitochondria was sufficient to upregulate protein production. Surprisingly, protein production was increased independent of the mRNA harboring an MTS, as an mRNA that contained *flag-GFP* with no mitochondrial coding sequences also showed increased protein production (Figures 6B and 6C). We then analyzed whether tethering to the ER might affect protein production by inserting the MCP protein into the C-terminus of Sec63p. We also saw increased protein production when mRNA was tethered to the ER, suggesting that the surface of both of these organelles may harbor the capacity for enhanced protein synthesis. In addition to increased protein expression, we also observed increased mRNA levels when mRNAs were tethered to the mitochondria. However, the ratio of protein to mRNA was much higher (Figure S10). We further observed a 3-fold increase in ribosome association rate from sucrose density fractionated mRNAs for mRNAs that were tethered to the mitochondria (Figure 6D), suggesting that translational efficiency is increased on the mitochondrial surface. To test whether localization to the mitochondria is necessary for optimal protein production during respiratory conditions, we reduced the localization of endogenous *ATP3* and *TIM50* mRNA to mitochondria by directing those mRNAs to the plasma membrane via insertion of a CaaX-tag to the C termini of MCP-GFP proteins during respiratory conditions (Yan et al., 2016). This caused a decrease in protein reporter levels of mitochondrial localized mRNAs, such as *TIM50, ATP3*, and *ATP2*, but not in non-mitochondrially localized mRNAs, such as *TOM22* and *GFP* alone (Figure 6E). We next investigated whether enhancing protein synthesis was essential for optimal cell growth. Cells in which *ATP3* mRNA was anchored to the plasma membrane and away from the mitochondria in respiratory conditions showed a growth defect, whereas ER tethering of mRNAs, which does not impair protein synthesis, did not affect cell growth (Figures 6F and S11). This suggests that localization of mRNA to mitochondria is important for optimal cell growth because it drives enhanced protein synthesis during respiratory conditions.

**Figure 6.**
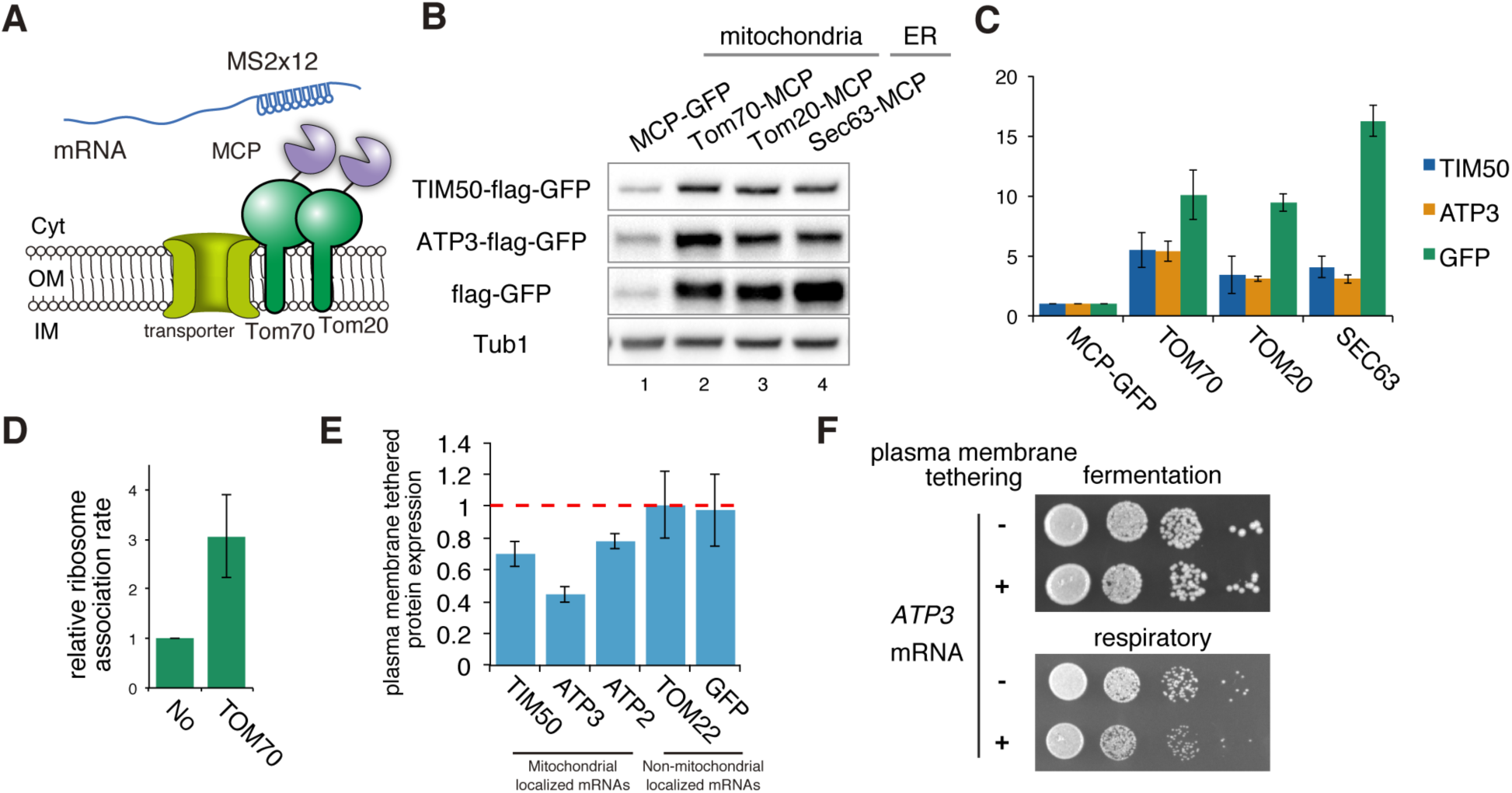
mRNA Localization to Mitochondria Enhances Its Translation. (A) Schematic of artificial mRNA tethering to mitochondria using MS2-MCP system. mRNAs harboring a MS2 tandem sequence were tethered to mitochondria through C-terminus MCP tagged Tom70p or Tom20p. (B) Protein expression analysis of reporter mRNAs, which are tethered to mitochondria (lanes 2, 3) or ER (lane 4). Protein expression was analyzed using an anti-GFP antibody. Tub1p in the strain harbouring GFP-MS2 reporter genes was used as an internal loading control. (C) Quantification of (B). Protein expression was normalized to the No-MCP strains. Error bar represents standard deviation of three independent experiments. (D) Relative ribosome association rate was calculated by comparison between RNA levels from ribosome-bound and free fractions from sucrose gradient polysome fractionation. RNA levels were quantified via qPCR. (E) Protein expression analysis in anchored-away conditions. Protein expression levels in each strain in respiratory conditions were determined by Western blotting using antiflag antibody. Protein expression ratio between strains with and without CaaX is shown as a bar graph. Error indicates standard deviation of three independent experiments. (F) Growth assay for plasma membrane localized mRNA, which consists of an integrated MS2 sequence into the 3’-UTR of genomic DNA. TIM50 and ATP3 mRNAs were anchored away to the plasma membrane using CaaX-tag harbored MCP-GFP proteins. Cell growth was tested on YPAD (fermentative) and YPAGE (respiratory) conditions at 30°C for 2 days and 3 days, respectively.

## Discussion

During fluctuating environmental conditions, cells must be able to control gene expression in order to optimize fitness. We demonstrate that yeast cells can use the geometric constraints that arise from increased mitochondrial volume fraction during respiratory conditions to drive condition-dependent mitochondrial localization for a subset of nuclear-encoded mitochondrial mRNAs. We favor the hypothesis that the geometric constraints of mitochondrial volume fraction are sufficient to explain our mRNA localization results for two reasons: first, our simple mathematical model incorporating mitochondrial volume fraction and various binding affinities is able to recapitulate the mRNA localization effects we see in cells. Second, the relationship between mitochondrial volume fraction and mRNA localization holds across a multitude of experimental perturbations, which presumably impact mitochondrial function in very different ways. *reg1*Δ changes mitochondrial volume fraction exclusively by increasing mitochondrial volume, whereas chloramphenicol and a nutrient shift from glucose to glycerol media increase mitochondrial volume fraction by both increasing mitochondrial volume and decreasing cytoplasmic volume. While *reg1*Δ and glycerol media increase oxidative phosphorylation, chloramphenicol inhibits mitochondrial translation and respiratory function. Finally, *sch9*Δ has a reduced mitochondrial volume but an increased vacuolar volume. This increase in vacuolar size decreases the accessible cytoplasm, thereby increasing the mitochondrial volume fraction, and increases *ATP3* mRNA localization to the mitochondria. While we believe the geometric constraints of the cell are import for mRNA localization, we cannot completely exclude the possibility that there are secondary factors that correlate with mitochondrial volume fraction and drive the mRNA localization effects we see because, to our knowledge, it is impossible to change mitochondrial volume fraction without perturbing multiple biological pathways.

Our results also indicate that translation elongation plays an important role in mRNA localization to the mitochondria. We believe that translation elongation and mitochondrial volume fraction are interconnected, as both will impact the probability of the mitochondria interacting with a competent mRNA, meaning the MTS is exposed while still attached to the mRNA as a nascent-polypeptide (Figure 7A). *ATP3* mRNA, which has fast translation elongation compared to *TIM50*, is in a competent state for a shorter period of time, so during fermentative conditions it is less likely to interact with mitochondria while in a competent state (Figure 7B). Conversely, ribosome stalling along *TIM50* mRNA prolongs the time the mRNA is in a competent state, and this increases its probability of interacting with mitochondria even when in low mitochondrial volume fraction conditions (Figure 7B). As the mitochondrial volume fraction of cells increases as they shift to respiratory conditions, there is a higher probability of competent mRNAs interacting with the mitochondria, even if they are only competent for a short time (Figure 7B).

**Figure 7.**
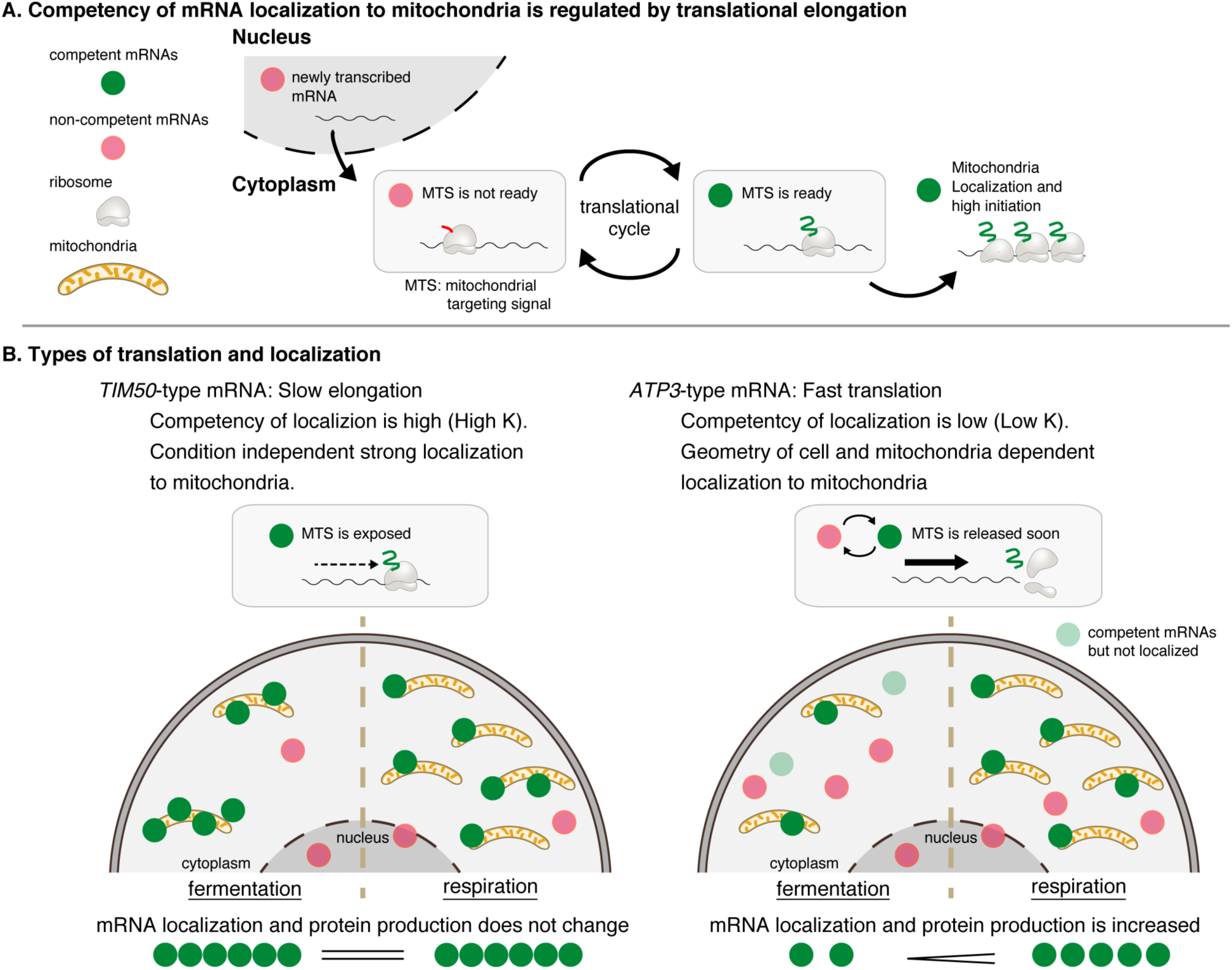
Mitochondrial Volume Fraction Controls Translation of Nuclear-Encoded Mitochondrial Proteins. (A) Competency of mRNA localization to mitochondria is regulated by translational elongation. After the MTS is translated and exposed from the ribosome the mRNA becomes competent for localization to the mitochondria for as long as the mRNA is associated with the nascent chain. Slow elongation will extend the time an mRNA is competent. Mitochondrially localized mRNAs experience high translation initiation. (B) Mitochondria can coordinate gene expression during times of metabolic need via mitochondrial-volume-fraction-based control and simple chemical kinetics of nuclearencoded mRNA localization. *TIM50*-type: mRNAs with high affinity to mitochondria are always associated with mitochondria and thus not much affected by geometrical features. *ATP3*-type: mRNAs with low affinity for mitochondria localization are greatly affected by geometrical features of cells, i.e., mitochondrial volume fraction. Fast translation elongation leads to quick release of the mRNA/nascent chain complex, and results in a quick return to a non-competent state. When mitochondrial volume fraction is high in respiratory conditions, mRNA localization to mitochondria is increased, and protein synthesis is induced by its localization.

The localization of ribosomes on the yeast mitochondrial surface has been known since the 1970s, implicating localized translation. It has been proposed that the functional relevance of this localized translation may be to facilitate co-translational import (Lesnik et al., 2015). The necessity of co-translational import is unclear, as *in vitro* mitochondrial import systems have found it to be important for the import of some proteins but not for others (Fujiki and Verner, 1991; Schatz, 1979; Suissa and Schatz, 1982). Our work points to localization to mitochondria having an alternative function: to upregulate protein synthesis, as we found that mRNA localization to mitochondria was sufficient to increase protein synthesis, whereas tethering mRNAs away from the mitochondria to the plasma membrane reduced protein synthesis. The decrease in protein levels when *ATP3* mRNA is anchored to the plasma membrane is also associated with a growth defect during respiratory conditions. Yet localization to the mitochondria is not absolutely required for optimal growth; when *ATP3* mRNAs were targeted away from the mitochondria to the ER, there was no growth defect. This may be because, similar to the mitochondria, there is an upregulation of protein levels when mRNAs are targeted to the ER.

Condition-dependent mRNA localization to the mitochondria as a means to control gene expression could be used to tune the protein composition of the mitochondria to the metabolic needs of the cell. Several observations support this hypothesis. For example, the conserved subunits of ATP synthase are all sensitive to the translation elongation rates in the cell, and these subunits showed similar localization regulation patterns (Table S3); this suggests that a mechanism may have evolved that coordinates the expression and stoichiometry of vital subunits of this complex. Further supporting this is global data showing that cycloheximide-sensitive mRNAs (*ATP3*-type) show a more than 2-fold increase in protein synthesis relative to constitutively localized mRNAs (*TIM50*-type) when cells shift from fermentation to respiration. For *ATP3*-type mRNAs, the increase in protein synthesis during respiratory conditions is driven by both increases in translational efficiency and increased mRNA levels (Figure 1C to 1E). Similarly, the increase in protein levels seen when mRNAs were tethered to the mitochondria was driven by both an increase in mRNA levels and ribosome engagement (Figures S9A and S9B). As these reporter mRNAs are under identical promoters, we favor the hypothesis that localization to the mitochondria increases mRNA stability. It is unclear whether translational efficiency and mRNA stability are independently affected by localization to the mitochondria or whether they are interrelated, as previous reports have shown that increased translation initiation can lead to protection from mRNA degradation (Roy and Jacobson, 2013). As discussed previously, ribosome stalling increases mRNA binding competency, driving mRNA localization to the mitochondria, along with increased translation initiation and mRNA stability. As translation initiation is thought to be the rate-limiting step in translation for most mRNAs, this would explain our somewhat-contradictory results showing that more ribosome stalling, through polyproline insertion, leads to increased protein synthesis.

How mitochondrially localized mRNAs can increase protein synthesis is still an open question. We consider the simplest explanation, that there may be an increased density of ribosomes on the mitochondrial surface, as previously suggested in *Drosophila* oogenesis (Zhang et al., 2016). It also might be true that translation initiation factors are highly phosphorylated around mitochondria, as mitochondria produce high levels of ATP. An alternative idea is that there are specialized ribosomes, enriched at the mitochondrial surface, that enhance translation (Xue and Barna, 2012). Intriguingly, it was recently found that specific ribosomal protein paralogs are necessary for normal mitochondrial function. *Rpl1bΔ* cells were found to be deficient for growth on nonfermentable carbon sources, whereas *rpl1aΔ* had no growth defect in these conditions. Furthermore, *rpl1bΔ* cells were found to have decreased translation of many mitochondrial proteins during respiratory conditions, including the *ATP3*-type mRNAs *ATP1* and *ATP2* (Segev and Gerst, 2018). Determining how mRNA localization increases protein synthesis will increase our understanding on how mitochondria are able to control their composition in relation to the metabolic needs of the cell.

While we have found this gene expression control mechanism in yeast, we speculate that higher eukaryotic cells could also use mitochondrial mRNA localization, controlled by translation elongation and mitochondrial volume fraction, to regulate protein synthesis. Hinting at this, a recent paper exploring global mRNA localization in mammalian cells found that many ATP synthase mRNAs had enhanced localization to the mitochondria in a cycloheximide-dependent manner, similar to *ATP3*-type mRNAs in yeast (Fazal et al., 2019). Further exploration into the role of mitochondrial volume fraction and translation elongation in mRNA localization to the mitochondria will provide insight into the regulation of mitochondria in health and disease.

## Methods

### Reconstruction of 3D Mitochondria and mRNA Visualization

To allow accurate visualization of mRNA molecules, multiple MS2 stem-loops are inserted in the 3’-UTR of the mRNA of interest and are recognized by the MCP-GFP fusion protein (Bertrand et al., 1998; Haim-Vilmovsky and Gerst, 2009). We improved this system by titrating down the MCP-GFP levels until we observed single molecule mRNA foci, which we verified by single molecule RNA FISH (smFISH) (Hocine et al., 2012; Tutucci et al., 2018) (Figure S1). We then performed rapid 3D live cell imaging using spinning disk confocal microscopy. We reconstructed and analyzed the spatial relationship between the mRNAs and mitochondria using custom ImageJ plugin Trackmate (Tinevez et al., 2017) and MitoGraph V2.0, which we previously developed to reconstruct 3D mitochondria based on matrix marker fluorescent protein intensity (Rafelski et al., 2012; Viana et al., 2015) (Figure 1A). We measured the distance between mRNA and mitochondria by finding the closest meshed surface area of the mitochondria matrix. Bias-reduced logistic regression (Firth, 1992, 1993) was used to determine which factors influenced the manual tracking of foci in Trackmate. Signal-to-noise ratio (SNR), median intensity of foci, and minimum distance between tracked foci were screened for their contributions to manual tracking of foci. Two-sided p-values were compared for the different variables. The logistic regression analysis shows that median intensity (p-value = 0.046) and SNR (p-value = 0.036) are the two features detectable by human eyes to track the foci.

### Definition of “Localization” and “Association” to Mitochondria

To analyze the association of mRNA to mitochondria, we first defined the “localized” threshold as twice the size of the mode (190nm) of the distance between *TIM50* mRNA and mitochondria, which were treated with translation elongation and transcription inhibitors cycloheximide (CHX) and 1,10-Phenanthroline (PHE), respectively, expecting that the majority of *TIM50* mRNAs associate with the mitochondrial surface in these conditions (Figure S2). We further classified mRNA localization to reflect a stable or transient association using this 0.19um threshold. Since the translational elongation rate is 9.5aa/sec (Shah et al., 2013), which implies that it takes more than 30sec to translate reporter genes (more than 300aa), we defined a mitochondria-associated mRNA as being co-localized to the mitochondria for at least 3 seconds (two consecutive time points).

## Supporting information

Tsuboi et al., 2019 bioRxiv_Supplementary Material

## SUPPLEMENTAL INFORMATION

Supplemental Information includes Extended Experimental Procedures, 11 figures, and 3 tables.

## ACKNOWLEDGMENTS

We thank members of the Zid laboratory as well as T. Endo, S. Iwasaki, G. Goshima, E. Koslover, W. Wang and V. Bilanchone for helpful discussions and A. Subramaniam and S. Mukherji for feedback on the paper. Receipt of the MS2-tagging plasmids and yeast-optimized fluorophore plasmids from Dr. J. Gerst and Dr. K. Thorn is gratefully acknowledged. We thank M. Zid, A. Guzikowski and V. Harjono for critically reading the manuscript. This work was supported in part by National Institutes of Health GM57071 (to R.H.S.), startup funds from UCI, NSF grant MCB-1330451 and Ellison Medical Foundation (to S.M.R), startup funds from UCSD and from the National Institutes of Health R35GM128798 (to B.M.Z.), and NINDS P30NS047101 (to UCSD microscopy Core). T.T. acknowledges support from a Japan Society for the Promotion of Science (JSPS) for a research abroad fellowship and postdoctoral fellowship (18J00995), and Uehara Memorial Foundation for research abroad fellowship.

## AUTHOR CONTRIBUTIONS

T.T., S.M.R., and B.M.Z. designed the study. T.T, and F.X. performed experiments. T.T., M.P.Z., J.Y., R.C., X.G.A., and Y.S.C. performed analysis. E.T., J.C. and R.H.S. advised on the research. T.T. and B.M.Z. wrote the manuscript. All authors discussed the results and commented on the manuscript.

## DATA AND MATERIALS AVAILABILITY

Further information and requests for resources, scripts, and reagents should be directed to and will be fulfilled by the lead contact, T.T.

## Bibliography

1. Adachi, A., Koizumi, M., and Ohsumi, Y. (2017). Autophagy induction under carbon starvation conditions is negatively regulated by carbon catabolite repression. J. Biol. Chem. 292, 19905–19918.

2. Arganda-Carreras, I., Kaynig, V., Rueden, C., Eliceiri, K.W., Schindelin, J., Cardona, A., and Sebastian Seung, H. (2017). Trainable Weka Segmentation: a machine learning tool for microscopy pixel classification. Bioinformatics 33, 2424–2426.

3. Attardi, G., and Schatz, G. (1988). BIOGENESIS OF MITOCHONDRIA. Ann. Rev. Cell Biol. 4, 289–333.

4. Bertrand, E., Chartrand, P., Schaefer, M., Shenoy, S.M., Singer, R.H., and Long, R.M. (1998). Localization of ASH1 mRNA Particles in Living Yeast. Mol. Cell 2, 437–445.

5. Borst, P., and Grivell, L.A. (1978). The Mitochondrial Genome of Yeast. Cell 15, 705–723.

6. Brody, S. (1992). Circadian rhythms in neurospora crassa: The role of mitochondria. Chronobiol. Int.

7. Chartron, J.W., Hunt, K.C.L., and Frydman, J. (2016). Cotranslational signal-independent SRP preloading during membrane targeting. Nature 536, 224–228.

8. Couvillion, M.T., Soto, I.C., Shipkovenska, G., and Churchman, L.S. (2016). Synchronized mitochondrial and cytosolic translation programs. Nature 533, 499–503.

9. De Deken, R.H. (1966). The Crabtree Effect: A Regulatory System in Yeast. J. Gen. Microbiol. 44, 149–156.

10. Egner, A., Jakobs, S., and Hell, S.W. (2002). Fast 100-nm resolution three-dimensional microscope reveals structural plasticity of mitochondria in live yeast. Proc. Natl. Acad. Sci. U. S. A. 99, 3370–3375.

11. Eliyahu, E., Pnueli, L., Melamed, D., Scherrer, T., Gerber, A.P., Pines, O., Rapaport, D., and Arava, Y. (2010). Tom20 Mediates Localization of mRNAs to Mitochondria in a Translation-Dependent Manner. Mol. Cell. Biol. 30, 284–294.

12. Elstner, M., Andreoli, C., Klopstock, T., Meitinger, T., and Prokisch, H. (2009). Chapter 1 The Mitochondrial Proteome Database. MitoP2. Methods Enzymol. 457, 3–20.

13. Fazal, F.M., Han, S., Parker, K.R., Kaewsapsak, P., Xu, J., Boettiger, A.N., Chang, H.Y., and Ting, A.Y. (2019). Atlas of Subcellular RNA Localization Revealed by APEX-Seq. Cell 1–18.

14. Firth, D. (1992). Bias reduction, the Jeffreys prior and GLIM. In Advances in GLIM and Statistical Modelling, L. Fahrmeir, B. Francis, R. Gilchrist, and G. Tutz, eds. (New York, NY: Springer New York), pp. 91–100.

15. Firth, D. (1993). Bias Reduction of Maximum Likelihood. Biometrika.

16. Fujiki, M., and Verner, K. (1991). Coupling of protein synthesis and mitochondrial import in a homologous yeast in vitro system. J. Biol. Chem. 266, 6841–6847.

17. Gadir, N., Haim-Vilmovsky, L., Kraut-Cohen, J., and Gerst, J.E. (2011). Localization of mRNAs coding for mitochondrial proteins in the yeast Saccharomyces cerevisiae. RNA 17, 1551–1565.

18. Garcia, M., Darzacq, X., Delaveau, T., Jourdren, L., Singer, R.H., and Jacq, C. (2007). Mitochondria-associated yeast mRNAs and the biogenesis of molecular complexes. Mol. Biol. Cell 18, 362–368.

19. Garcia, M., Delaveau, T., Goussard, S., and Jacq, C. (2010). Mitochondrial presequence and open reading frame mediate asymmetric localization of messenger RNA. EMBO Rep. 11, 285–291.

20. Gleason, F.K., Ooka, M.P., Cunningham, W.P., and Hooper, A.B. (1975). Effect of chloramphenicol on replication of mitochondria in Tetrahymena. J. Cell. Physiol.

21. Golani-Armon, A., and Arava, Y. (2016). Localization of Nuclear-Encoded mRNAs to Mitochondria Outer Surface. Biochemistry. (Mosc). 81, 1038–1043.

22. Gold, V.A., Chroscicki, P., Bragoszewski, P., and Chacinska, A. (2017). Visualization of cytosolic ribosomes on the surface of mitochondria by electron cryo-tomography. EMBO Rep. e201744261.

23. Haim-Vilmovsky, L., and Gerst, J.E. (2009). m-TAG: a PCR-based genomic integration method to visualize the localization of specific endogenous mRNAs in vivo in yeast. Nat. Protoc. 4, 1274–1284.

24. Hocine, S., Raymond, P., Zenklusen, D., Chao, J.A., and Singer, R.H. (2012). Single-molecule analysis of gene expression using two-color RNA labeling in live yeast. Nat. Methods 10, 119–121.

25. Hubscher, V., Mudholkar, K., Chiabudini, M., Fitzke, E., Wolfle, T., Pfeifer, D., Drepper, F., Warscheid, B., and Rospert, S. (2016). The Hsp70 homolog Ssb and the 14-3-3 protein Bmh1 jointly regulate transcription of glucose repressed genes in Saccharomyces cerevisiae. Nucleic Acids Res. 44, 5629–5645.

26. Jorgensen, P., Nishikawa, J.L., Breitkreutz, B.-J., and Tyers, M. (2002). Systematic Identification of Pathways That Couple Cell Growth and Division in Yeast. Science 297, 1070850.

27. Kellems, R.E., Allison, V.F., and Butow, R.A. (1974). Cytoplasmic Type 80 S Ribosomes Associated with Yeast Mitochondria. II. EVIDENCE FOR THE ASSOCIATION OF CYTOPLASMIC RIBOSOMES WITH THE OUTER MITOCHONDRIAL MEMBRANE IN SITU. J. Biol. Chem. 249, 3297–3303.

28. Lesnik, C., Cohen, Y., Atir-Lande, A., Schuldiner, M., and Arava, Y. (2014). OM14 is a mitochondrial receptor for cytosolic ribosomes that supports co-translational import into mitochondria. Nat. Commun. 5, 1–10.

29. Lesnik, C., Golani-armon, A., and Arava, Y. (2015). Localized translation near the mitochondrial outer membrane : An update. RNA Biol. 12, 801–809.

30. Marc, P., Margeot, A., Devaux, F., Blugeon, C., Corral-Debrinski, M., and Jacq, C. (2002). Genome-wide analysis of mRNAs targeted to yeast mitochondria. EMBO Rep. 3, 159–164.

31. Martin, K.C., and Ephrussi, A. (2009). mRNA Localization: Gene Expression in the Spatial Dimension. Cell 136, 719–730.

32. Morgenstern, M., Stiller, S.B., Lübbert, P., Peikert, C.D., Dannenmaier, S., Drepper, F., Weill, U., Höß, P., Feuerstein, R., Gebert, M., et al. (2017). Definition of a High-Confidence Mitochondrial Proteome at Quantitative Scale. Cell Rep. 19, 2836–2852.

33. Paulo, J.A., O’Connell, J.D., Everley, R.A., O’Brien, J., Gygi, M.A., and Gygi, S.P. (2016). Quantitative mass spectrometry-based multiplexing compares the abundance of 5000 S. cerevisiae proteins across 10 carbon sources. J. Proteomics 148, 85–93.

34. Rafelski, S.M., Viana, M.P., Zhang, Y., Chan, Y.-H.M., Thorn, K.S., Yam, P., Fung, J.C., Li, H., Costa, L.D.F., and Marshall, W.F. (2012). Mitochondrial network size scaling in budding yeast. Science 338, 822–824.

35. Reid, D.W., and Nicchitta, C. V. (2015). Diversity and selectivity in mRNA translation on the endoplasmic reticulum. Nat. Rev. Mol. Cell Biol. 16, 221–231.

36. Roy, B., and Jacobson, A. (2013). The intimate relationships of mRNA decay and translation. Trends Genet.

37. Saint-Georges, Y., Garcia, M., Delaveau, T., Jourdren, L., Le Crom, S., Lemoine, S., Tanty, V., Devaux, F., and Jacq, C. (2008). Yeast mitochondrial biogenesis: a role for the PUF RNA-binding protein Puf3p in mRNA localization. PLoS One 3, e2293.

38. Schatz, G. (1979). How mitochondria import proteins from the cytoplasm. FEBS Lett.

39. Segev, N., and Gerst, J.E. (2018). Specialized ribosomes and specific ribosomal protein paralogs control translation of mitochondrial proteins. J. Cell Biol. 217, 117–126.

40. Shah, P., Ding, Y., Niemczyk, M., Kudla, G., and Plotkin, J.B. (2013). Rate-limiting steps in yeast protein translation. Cell 153, 1589–1601.

41. Suissa, M., and Schatz, G. (1982). Import of Proteins into Mitochondria. J. Biol. Chem. 257, 13048–13055.

42. Tinevez, J.Y., Perry, N., Schindelin, J., Hoopes, G.M., Reynolds, G.D., Laplantine, E., Bednarek, S.Y., Shorte, S.L., and Eliceiri, K.W. (2017). TrackMate: An open and extensible platform for single-particle tracking. Methods.

43. Tutucci, E., Vera, M., Biswas, J., Garcia, J., Parker, R., and Singer, R.H. (2018). An improved MS2 system for accurate reporting of the mRNA life cycle. Nat. Methods 15, 81–89.

44. Viana, M.P., Lim, S., and Rafelski, S.M. (2015). Quantifying mitochondrial content in living cells (Elsevier Ltd).

45. Williams, C.C., Jan, C.H., and Weissman, J.S. (2014). Targeting and plasticity of mitochondrial proteins revealed by proximity-specific ribosome profiling. Science (80-.). 346, 748–751.

46. Williamson, D.H., Maroudas, N.G., and Wilkie, D. (1971). Induction of the cytoplasmic petite mutation in Saccharomyces cerevisiae by the antibacterial antibiotics erythromycin and chloramphenicol. MGG Mol. Gen. Genet. 111, 209–223.

47. Wu, B., Eliscovich, C., Yoon, Y.J., and Singer, R.H. (2016). Translation dynamics of single mRNAs in live cells and neurons. Science (80-.). 352, 1430–1435.

48. Xue, S., and Barna, M. (2012). Specialized ribosomes: A new frontier in gene regulation and organismal biology. Nat. Rev. Mol. Cell Biol. 13, 355–369.

49. Yan, X., Hoek, T.A., Vale, R.D., and Tanenbaum, M.E. (2016). Dynamics of Translation of Single mRNA Molecules in Vivo. Cell 165, 976–989.

50. Zhang, Y., Chen, Y., Gucek, M., and Xu, H. (2016). The mitochondrial outer membrane protein MDI promotes local protein synthesis and mtDNA replication. EMBO J.

51. Zid, B.M., and O’Shea, E.K. (2014). Promoter sequences direct cytoplasmic localization and translation of mRNAs during starvation in yeast. Nature 514, 117–121.

