## Supplementary material for "Mitochondrial volume fraction and translation speed impact mRNA localization and production of nuclear-encoded mitochondrial proteins": Tsuboi et al., 2019 bioRxiv_Supplementary Material

**This PDF file includes:**

EXPERIMENTAL PROCEDURES

Figures S1 to S11

Tables S1 to S3

References

### EXPERIMENTAL PROCEDURES

#### Yeast Strains and Plasmids

The yeast strains and plasmids used are listed in Supplementary Table S1 and the oligonucleotides used for plasmid construction are described in Table S2. To reduce variability among the constructed yeast strains, the strains were created either through integration of a linear PCR product or a plasmid linearized through restriction digest. Yeast strains harboring the genomic MS2 sequence were constructed using pSH47 and pLOXHIS5MS2L, a gift from the J. Gerst laboratory, as previously described (Gadir et al., 2011; Haim-Vilmovsky and Gerst, 2009). To constitutively visualize mRNAs, pRS405*CYC1p-MS2-4xGFP* (TTP080) was constructed as follows: *MCP-4xGFP* sequence was PCR amplified from pMS2CPGFP(x4) (Gadir et al., 2011) and integrated into the pRS405 vector under the *CYC1* promoter. The strains expressing MCP-4xGFP from 2 sets of integrated plasmids were selected through microscopy screening and used for further experiments. To visualize the mitochondria matrix, pRS406GPDp-Su9-mCherry was constructed as follows: Su9 (subunit 9 of the $F_0$ ATPase (Su9(1-69))) sequence was PCR amplified from pvt100-dsRed (Rafelski et al., 2012) and the yomCherry sequence was PCR amplified from pFA6a-link-yomCherry-SpHis5 (Lee et al., 2013), a gift from the K. Thorn laboratory, and integrated into pRS406 vector under a *GPD* promoter. The strains expressing Su9-mCherry from two sets of integrated plasmids were selected through microscopy screening. To construct plasmids shown in Figure 3A (TTP134 and TTP136), endogenously MS2 sequence-tagged genes were amplified from genomic DNA of strains TTY1160 and TTY439, and inserted into pRS403 vector plasmids (TTP133 and TTP135), respectively. Annealed cytERM (C450_2C1) primers were integrated into TTP133 and TTP135 by combination of Gibson assembly and PCR. GFP tagging was performed with PCR mediated homologous recombination using pFA6a-link-yoGFP-SpHis5 (Lee et al., 2013) and integrations were confirmed by PCR. Deletion mutant strains were constructed with PCR mediated homologous recombination using pFA6a-hphMX6, a gift from the J. Wilhelm laboratory, and integrations were confirmed by PCR. To construct the plasmids used in Figure 2 and 3, a yoGFP fragment was PCR-amplified from pFA6a-link-yoGFP-SpHis5 (Lee et al., 2013) or an iRFP fragment was PCR-amplified from NHB084 with a flag sequence and inserted into TTP133 to construct TTP155 and TTP146, respectively. Synthetic reporters were created by Gibson Assembly of a PCR-amplified *TIM50* (1-300nt), *TIM50* (301-1431nt), *ATP3* (1-300nt) and/or *ATP3* (301-936nt). To construct the polyproline deletion constructs TTP174 and TTP179, deletions were introduced by PCR-based site-directed mutagenesis. To endogenously insert MCP proteins into the C-terminus of Tom70p, Tom20p, Sec63p, we generated TTP119 and TTP153 by replacing a PCR amplified MCP fragment from pMS2CPGFP(x4) (Gadir et al., 2011) with yoGFP of pFA6a-link-yoGFP-SpHis5 and pFA6a-link-yoGFP-CaUra3 (Lee et al., 2013). MCP tagging was performed with PCR mediated homologous recombination using TTP119 and TTP153. Integrations were confirmed by PCR followed by sequencing. Plasma membrane anchor MCP-CaaX was constructed by PCR-based site-directed mutagenesis using TTP080. The strains expressing MCP-2xGFP-CaaX from two sets of integrated plasmids were selected through microscopy screening and used for further experiments.

**Microscopy**

Single molecule mRNA visualization with mitochondria was performed as follows: 300μl of mid-log phase wild-type yeast cells (OD600 of 0.4-0.7), grown in appropriate medium, were harvested and placed into an Y04C microfluidic chamber, controlled by the CellASIC Onix system. 300µl YPAD were placed into the flow-wells, the chambers were loaded with cells at 3psi, and medium was continuously flowed at 3psi. Cells were imaged at 30°C with a Yokogawa CSUX Spinning Disk Confocal (Solamere Technology Group) mounted on a Nikon Eclipse Ti chassis motorized inverted microscope, located at the Department of Developmental & Cell Biology (UCI). Imaging was performed using a 100x/1.49 NA oil APO TIRF objective with the correction collar set manually for each experiment and a 1x tube lens (pixel size 0.084µm). Z-stacks (300nm steps) were acquired in the fluorescent channel (33ms exposure) on a Hamamatsu electron-multiplying charge-coupled device (EMCCD) camera. Imaging was controlled using MicroManager ImageAcquisition (v1.4.16). For CHX treatment, a flow-well containing a final concentration of 100µg/ml Cycloheximide (C7698; Sigma-Aldrich) was open and an image was taken every 3sec. To directly analyze the effect of *bona fide* inhibition of translation, we supplemented 1,10-Phenanthroline (131377; Sigma-Aldrich), which inhibits transcription, to CHX with final concentration of 250µg/ml. For LTM treatment, a flow-well containing a final concentration of 50µM Lactimidomycin (506291; MilliporeSigma) was open for 20min and an image was taken every 3sec. Image data for Figure 2E and S5 as well as C-terminus integrated GFP intensity data were collected as follows: Mid-log phase wild-type yeast cells (OD600 of 0.4 to 0.7) were grown in appropriate medium and 100µl were placed into a 96-well Glass Bottom Plate (Cellvis LLC). For Chloramphenicol treatment, cells were grown in YPAD medium containing a final concentration of 1mg/ml Chloramphenicol (C0378; Sigma-Aldrich). Cells were imaged at 23°C with a Perkin Elmer UltraView Vox Spinning Disk Confocal mounted on an Olympus IX81 inverted microscope with Yokogawa CSU-X1-A1 spinning disk head, located at the UCSD School of Medicine Microscopy Core. Imaging was performed using either 100x/1.4 NA or 60x/1.42 NA oil objective with the correction collar set manually for each experiment (pixel size 0.68µm or 0.111µm). Z-stacks (300nm steps) were acquired in the fluorescent channel on an EMCCD Hamamatsu 14 bit 1K x 1K camera. Imaging was controlled using Volocity (PerkinElmer).

**Reconstruction of 3D Mitochondria and mRNA Visualization**

To allow accurate visualization of mRNA molecules, multiple MS2 stem-loops are inserted in the 3'-UTR of the mRNA of interest and are recognized by the MCP-GFP fusion protein (Bertrand et al., 1998; Haim-Vilmovsky and Gerst, 2009). We improved this system by titrating down the MCP-GFP levels until we observed single molecule mRNA foci, which we verified by single molecule RNA FISH (smFISH) (Hocine et al., 2012; Tutucci et al., 2018a) (Figure S1). We then performed rapid 3D live cell imaging using spinning disk confocal microscopy. We reconstructed and analyzed the spatial relationship between the mRNAs and mitochondria using custom ImageJ plugin Trackmate (Tinevez et al., 2017) and MitoGraph V2.0, which we previously developed to reconstruct 3D mitochondria based on matrix marker fluorescent protein intensity (Rafelski et al., 2012; Viana et al., 2015) (Figure 1A). We measured the distance between mRNA and mitochondria by finding the closest meshed surface area of the mitochondria matrix. Bias-reduced logistic regression (Firth, 1992, 1993) was used to determine which factors influenced the manual tracking of foci in Trackmate. Signal-to-noise ratio (SNR), median intensity of foci, and minimum distance between tracked foci were screened for their contributions to manual tracking of foci. Two-sided p-values were compared for the different variables. The logistic regression analysis shows that median intensity (p-value = 0.046) and SNR (p-value = 0.036) are the two features detectable by human eyes to track the foci.

**Validation of Single Molecule mRNA of MS2-MCP System**

It has been observed that version 4 of the MS2-MCP system is prone to cytoplasmic aggregation, thought to be caused by either stabilization of mRNA or its degradation intermediates by MCP proteins (Tutucci et al., 2018a). We improved the MS2-MCP system by titrating down the MCP-GFP levels until we could observe single molecule mRNA foci, which we verified by single molecule RNA FISH (smFISH) (Hocine et al., 2012) (Figure S1). Analysing the co-localization rate of candidate mRNA ORF and MS2 sequences using anti-sense probes (Figure S1A), we confirmed that more than 84% of the ORFs and MS2 sequences are a continuous stretch in the same mRNA (Figure S1B). The number of MCP-GFP foci in live cells is statistically no different (Wilcoxon rank-sum test, p-value<0.05, N>43) than the number of smFISH foci in the tested mRNAs (Figure S1C). We concluded that the MCP-GFP foci in the live cell images are reliable single molecule mRNAs and not degradation intermediates.

**Single-Molecule FISH and Image Acquisition and Analysis**

Single-molecule FISH (smFISH) was performed as previously described (Tutucci et al., 2018b). Yeast strains were grown at 23°C in YPAD medium containing 2% glucose and harvested at OD600 0.6–0.8. Cells were fixed by adding paraformaldehyde (15714; 32% solution, EM grade; Electron Microscopy Science) to a final concentration of 4%. Cells walls were removed by resuspension in spheroplast buffer (buffer B containing 20mM VRC (S1402S ; Ribonucleoside–vanadyl complex NEB), and 25U of Lyticase enzyme (L2524; Sigma). Digested cells were seeded on 18mm polylysine-treated coverslips. The hybridization mix (10% formamide (205821000; ACROS organics), 2x SSC, 1mg/ml BSA, 10mM VRC, 5mM NaHPO4 pH 7.5, 1mg/µL E. coli tRNA, 1mg/µl ssDNA) was prepared with probe mix (final concentration

125nM). Cells were then hybridized at 37°C for 4hrs in the dark. Nuclei were stained with 0.5µg/mL DAPI in 1x PBS. Coverslips were mounted on glass slides using ProLong™ Gold Antifade Mountant (ThermoFisher). Images were acquired using an Olympus BX61 wide-field epifluorescence microscope with a 100x/1.35 NA UPlanApo objective. Samples were visualized using an X-Cite 120 PC lamp (EXFO) and the ORCA-R2 Digital CCD camera (Hamamatsu). Metamorph software (Molecular Devices) was used for acquisition. Z-sections were acquired at 200 nm intervals over an optical range of 8.0µm. Image pixel size: XY, 64.5nm. smFISH probes for *TIM50*, *ATP3* and *TOM22* probes were designed using and purchased from the Stellaris™ Probe Designer by LGC Biosearch Technologies. MS2 sequence probes, gifted from the R. Singer laboratory, were synthetized by Invitrogen-Thermo Fisher and labelled in the lab using Cy3 dyes (Amersham) as previously described (Tutucci et al., 2018b). Quantification of the FISH spots in the cytoplasm was performed using a custom ImageJ pipeline and manually detected. If the focus in different channels was within two pixels, it was considered to be co-localized; otherwise, it was considered to be two non-overlapping and distinct foci.

**Definition of "Localization" and "Association" to Mitochondria**

To analyze the association of mRNA to mitochondria, we first defined the "localized" threshold as twice the size of the mode (190nm) of the distance between *TIM50* mRNA and mitochondria, which were treated with translation elongation and transcription inhibitors cycloheximide (CHX) and 1,10-Phenanthroline (PHE), respectively, expecting that the majority of *TIM50* mRNAs associate with the mitochondrial surface in these conditions (Figure S2). We further classified mRNA localization to reflect a stable or transient association using this 0.19um threshold. Since the translational elongation rate is 9.5aa/sec (Shah et al., 2013), which implies that it takes more than 30sec to translate reporter genes (more than 300aa), we defined a mitochondria-associated mRNA as being co-localized to the mitochondria for at least 3 seconds (two consecutive time points).

**Segmentation of Cell Boundaries and Analysis of Cell Volume**

We segmented cell boundaries from the GFP field of the image, which we acquired for mRNA localization analysis in Figure 1B, using the ImageJ plug-in Trainable Weka Segmentation (Arganda-Carreras et al., 2017). Cell volume was calculated through this boundary. To analyze the effective mitochondrial volume fraction for analyzing mutant strains, we segmented the non-cytoplasmic area as vacuole. To acquire the volume fraction of the vacuole, we chose the center focal plane of each cell in bright field image and segmented the outer cell shape and inner black area, which we assumed as non-cytoplasmic area. Then, fit the ellipse to both outer cell shape and inner non-cytoplasmic area and calculate the volume using shorter radios of the fitted ellipse as depth.

**Single Molecule mRNA in silico Experiment and Mathematical Modeling**

We reconstructed the spatial boundaries of the cell and mitochondria in Paraview (http://www.paraview.org) and, to model ideal Brownian particles, we generated particle trajectories using a random walk in the space between the cell wall and mitochondria surface for at least 40,000 trajectories per cell. Then the mitochondrial localization ratio per cell (as we defined previously) was quantified. Because the particles were generated as space filling between the cell wall and mitochondrial surface and the mitochondria is a tubular structure, the mitochondrial localization ratio $R_{localized}$ was linearly correlated with mitochondria volume fraction as described in the following equation (Figure S4A, black line).

$$R_{localized} \propto \frac{mitochondrial\ surface\ area}{cell\ volume} \propto mitochondrial\ volume\ fraction$$

We assessed whether the localization behavior for *TOM22* mRNA, which we assumed to be a non-mitochondrial localized mRNA, matched with that of a Brownian particle. Although the mitochondrial localization ratio for *TOM22* mRNA was linearly correlated with the mitochondrial volume fraction, the slope was different from that of a Brownian particle. We hypothesized this was because we had not considered the other cell compartments, such as nucleus (7% of cell volume) (Jorgensen et al., 2007), vacuoles (10% of cell volume) (Chan et al., 2016), and mitochondria itself (10% of cell volume; this study). We determined that the difference in this ratio was caused by this organelle volume and to account for it, we minimized the cell size accordingly. To fit the *TOM22* mRNA localization ratio line to the Brownian particles we set the effective cell volume to 66%. This was below our expectation, but we found this acceptable since we had not included other organelles such as ER or lipid droplets in the virtual cell to study localization.

We assumed that mitochondrial localized mRNA concentration $[mRNA_{localized}]$, free diffusing mRNA concentration $[mRNA_{free}]$, and the concentration of mitochondrial surface point where mRNA can bind $[mito_{surface}]$ are in thermodynamical equilibrium. We then hypothesized that some mRNAs may associate with the mitochondrial surface by a sequence-specific association (e.g. via an MTS) with a constant K such that:

$$[mRNA_{free}][mito_{surface}] \rightleftarrows [mRNA_{localized}]$$

$$K = \frac{[mRNA_{localized}]}{[mito_{surface}][mRNA_{free}]}$$

$K$ is also thermodynamically given by

$$K = e^{-\Delta G_{localization}/_{k_B T}}$$

where $k_B$ is Boltzmann's constant, T is temperature, and $\Delta G_{localized}$ is the free energy gain from mitochondrial localization. For the Brownian particle-like *TOM22,* we assumed a $\Delta G_{localized}$ value of 0; therefore, *TOM22* had the lowest equilibrium constant, which we called $K_o$. We then hypothesized that any arbitrary mRNA has an equilibrium constant of $K=AK_o$, where A thus represents a fold-change in affinity between that mRNA and mitochondria compared to a freely diffusing mRNA. As the value of A increased in the simulation, the ratio of mitochondrial localization of the mRNA for a given mitochondrial volume fraction increased as well (Figure S4B). The mitochondrial localization ratio $R_{localized}'$ for an arbitrary mRNA when $K=AK_o$ is fitted to the following equation using the linear trend line for localization ratio ${R_{localized}}_0$ of *TOM22* mRNA.

$$R_{localized}' = \frac{A \cdot {mRNA_{localized}}_0}{1 + (A-1){R_{localized}}_0}$$

Thus, we show that the probability of an mRNA localizing to a mitochondrion is dependent upon both the mitochondrial concentration and the free energy change of localizing.

**Quantification of Protein Expression in Each Cell from Image Data**

Quantification of GFP-tagged mitochondrially-localized protein expression was performed using a custom analysis pipeline. The MitoGraph was applied to the mCherry channel to identify the locations of mitochondria and the resulting binary mask was then segmented to determine projections of individual mitochondria. Next, the sum of the intensity of the GFP channel covered by binary mask from each cell was measured using a custom ImageJ script.

**Western Blotting**

Yeast cells were grown in YPA medium containing 2% glucose or 3% glycerol + 2% ethanol. The cells were harvested when the culture reached an OD600 of 0.8. The protein products of various GFP reporter genes were detected by Western blotting using an anti-GFP antibody (11814460001; Roche) and a Mouse IgG (H+L) Secondary Antibody (32430; Thermo Fisher). For the detection of the flag-tagged proteins, anti-flag antibody (F1804; Sigma-Aldrich) was used. A mouse anti-a-tubulin antibody (12G10, Developmental Studies Hybridoma Bank) was used as a loading control. The intensities of the bands on the blots were quantified using Molecular Imager ChemiDoc XRS+ System (Bio-Rad) and ImageJ. The relative levels of the protein products were determined by comparison with a standard curve prepared using a series of dilutions.

**Polysome Analysis**

Yeast cells were grown exponentially at 30°C and harvested by centrifugation. Cell extracts were prepared with mortar and pestle as described previously (Tsuboi et al., 2012). The equivalent of 10 A260 units was then layered onto linear 10% to 50% (w/w) sucrose density gradients. Sucrose gradients (10%–50% sucrose in 20mM HEPES [pH 7.4], 5 mM magnesium chloride, 100 mM potassium chloride, 2 mM DTT, 100µg/ml CHX) were prepared in 14 x 89 mm polyallomer tubes (Beckman Coulter) with a gradient master. Crude extracts were layered on top of the sucrose gradients and were centrifuged at 30,000 rpm in a SW-28 rotor for 2.5 hr at 4°C . Gradients were fractionated using a gradient fractionator and UA-6 detector (ISCO/BRANDEL). RNA from ribosome-free and ribosome-bound fraction were purified from each fraction using guanidine hydrochloride and processed for RT-qPCR.

**RT-qPCR**

RNA was extracted using the MasterPure Yeast RNA Purification Kit (Epicentre). cDNA was prepared using ProtoScript® II Reverse Transcriptase (NEB #M0368X) with a combination of oligo(dT) primers and random hexamers according to the manufacturer's instructions. mRNA abundance was determined by qPCR using SYBR Green PCR Master Mix (Applied Biosystems) and primers specific for *FLAG-GFP* (ATGGACTACAAGGACGACG and CCTCACCACTCACAGAAAAC) and *ACT1* (GAGAGGCGAGTTTGGTTTCA and TCACCCGGCCTCTATTTTC). The mRNA levels were normalized to *ACT1* abundance, and the fold change between samples was calculated by a standard ΔΔCt analysis.

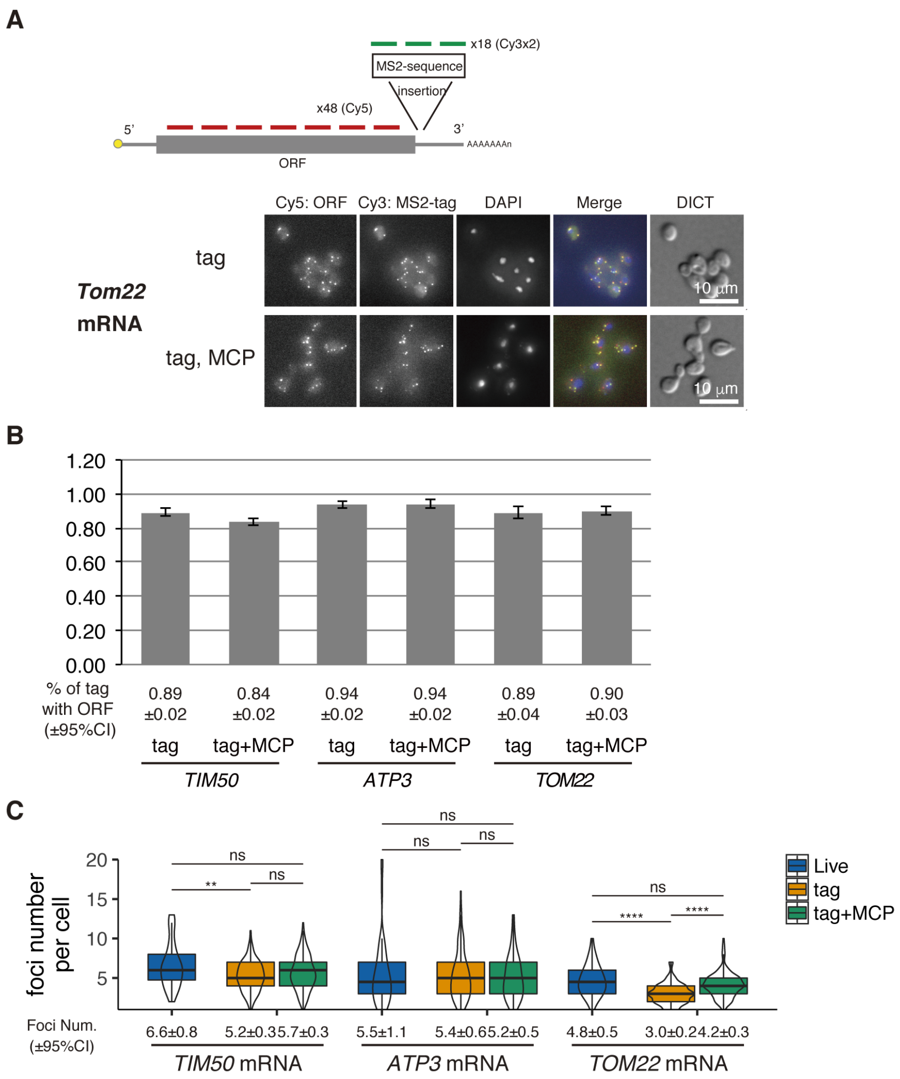


**Figure S1. Validation of Live Imaging mRNA Foci by Single Molecule FISH, Related to Figure 1**

(A) *TOM22*-tagged cells with or without CYC1p-MS2-CP-GFP(x4) were examined by a smFISH experiment. Fluorophore-conjugated multiple 20nt FISH probes were prepared to hybridize the ORF (Cy5) and MS2-tag (Cy3) as depicted.

(B) The ratio of MS2-tag with ORF. Percentages of the co-localized foci from the ORF and MS2-tag in MS2-tag foci were calculated based on manual count (N>103). Error bar represents s.e.m.

(C) Comparison between live imaging and FISH experiment foci number. The number of mRNA foci in live imaging and number of Cy3 Foci (MS2-tag) in FISH experiments were manually counted (N>43). Statistical significance was assessed by Mann–Whitney U-test (**** P < 0.0001; ** P < 0.01; ns, no significant difference).

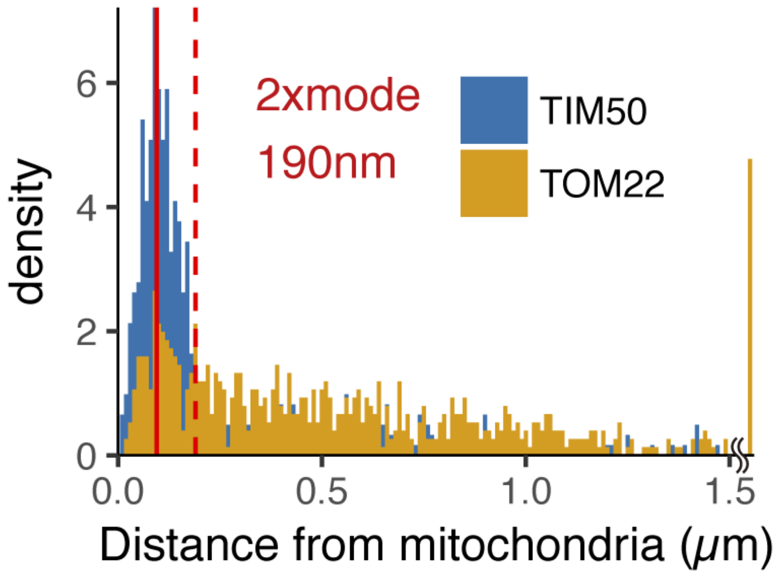


**Figure S2. The Distribution of mRNA-mitochondria Distance in Cycloheximide and 1,10-Phenanthroline Treated Cells, Related to Figure 1**

Number of foci in each mRNA is as follows: *TIM50* mRNA, 582; *TOM22* mRNA, 476. The red solid and dotted line indicate mode and 2x mode (0.19μm) of the distance between mitochondria and *TIM50* mRNA. The bin width is 0.01μm.

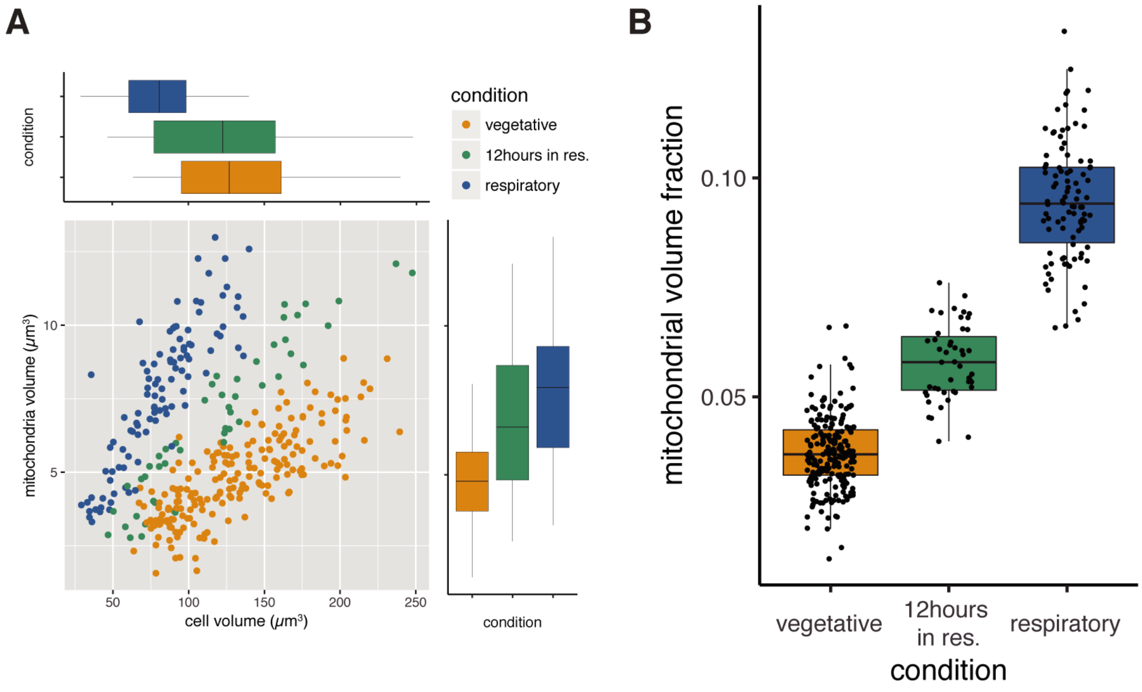


**Figure S3. Higher Mitochondrial Volume Fraction is a Feature of Respiratory Conditions, Related to Figure 1**

(A) Relationship of cell volume and mitochondrial volume of single cells in different growth conditions. "12 hours in res." corresponds to cells grown in respiratory medium for twelve hours after fermentative grown, while normal respiratory samples were grown in respiratory medium for the duration of the experiment.

(B) Mitochondrial volume fraction of single cells in different growth conditions.

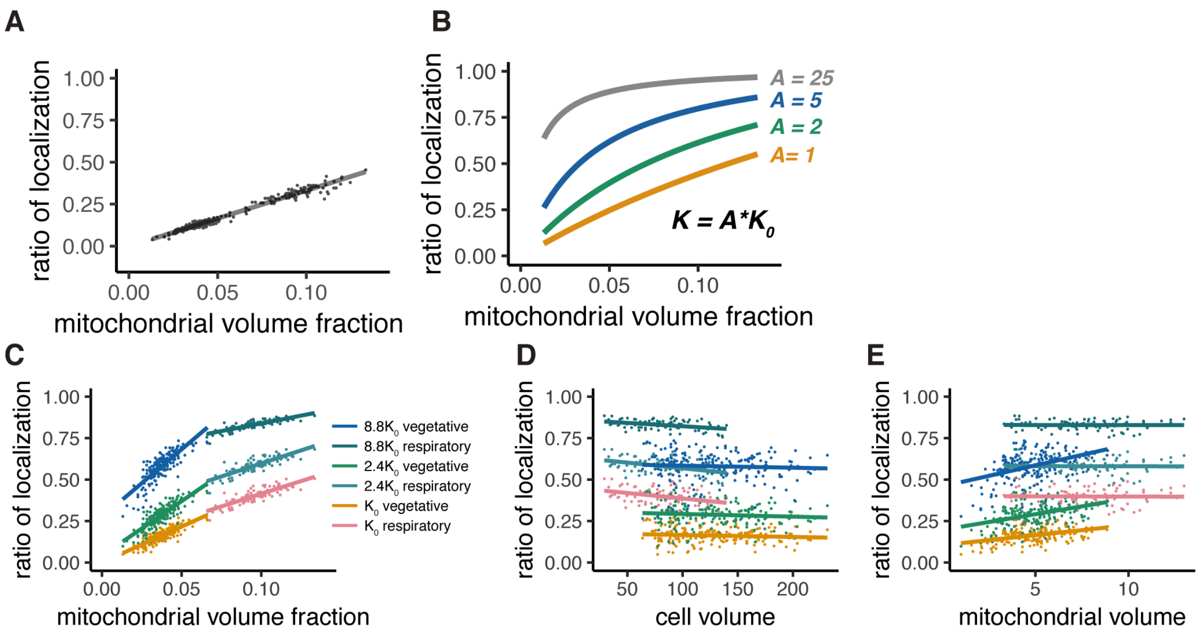


**Figure S4. Mitochondrial Volume Fraction and mRNA Localization Have Stoichiometric Correlation, Related to Figure 2**

(A) Relationship of ratio of mRNA localization to mitochondria and mitochondrial volume fraction from *in silico* experiments as described in Methods.

(B) Relationship of ratio of mRNA localization to mitochondria and mitochondrial volume fraction acquired by thermodynamically defined A values. The green, blue, and gray lines were plotted through equilibrium constant of A = $2K_0$, $5K_0$, and $25K_0$, respectively, as described in Methods.

(C) Mitochondrial volume fraction and mRNA localization have a stoichiometric correlation. Linear regression lines of the ratio of mRNA localization to mitochondria and mitochondrial volume fraction were plotted. Each line represents a linearly fitted line for $K_0$, $2.4K_0$, and $8.8K_0$ in fermentative and respiratory conditions, respectively.

(D) Cell volume and mRNA localization do not have a stoichiometric correlation. The same data as C was summarized with cell volume as the x-axis.

(E) Mitochondrial volume and mRNA localization do not have a stoichiometric correlation. The same data as C was summarized with mitochondrial volume as the x-axis.

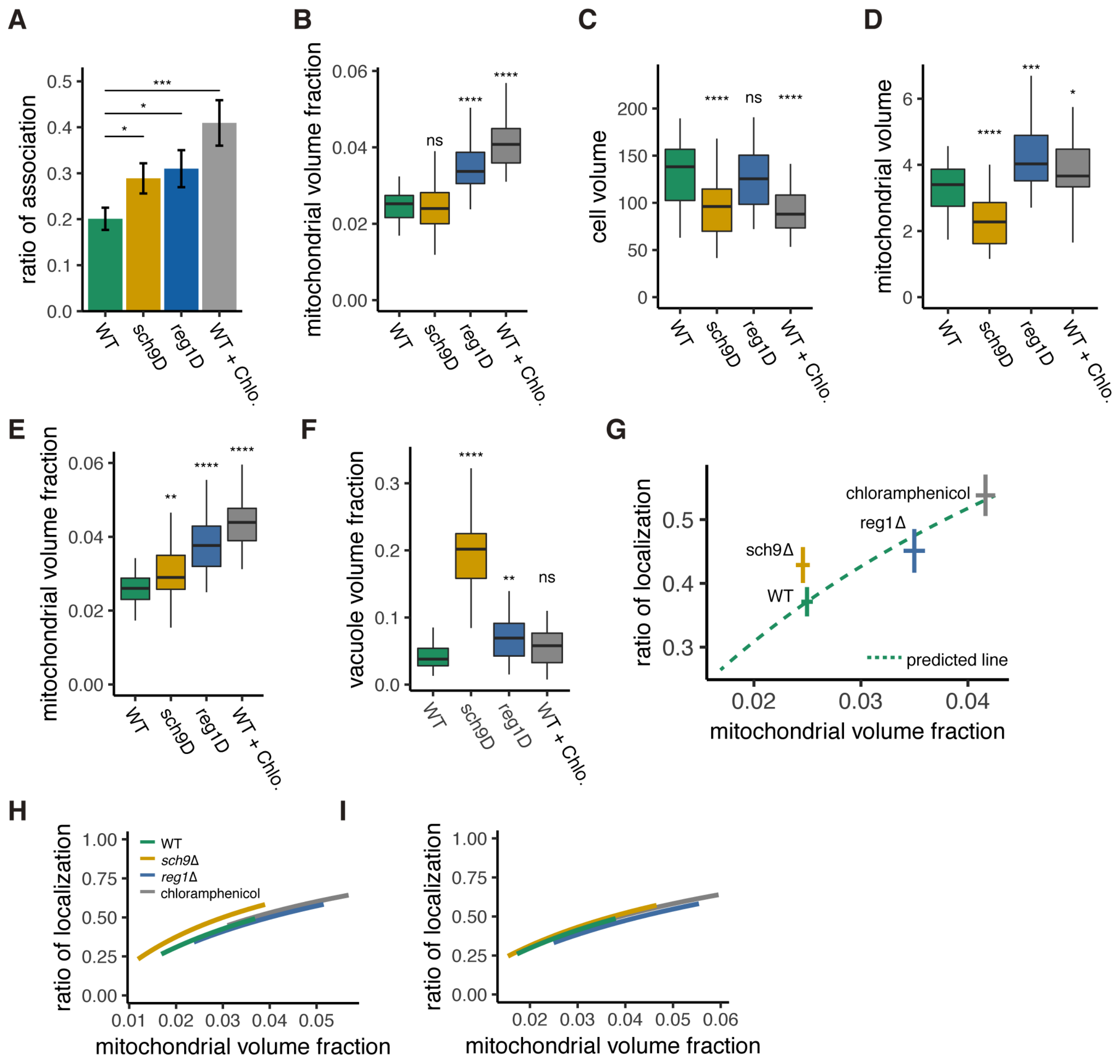


**Figure S5. Mitochondrial Volume Fraction Correlates with *ATP3* mRNA Localization in WT, *sch9*Δ and *reg1*Δ Mutant Strains, and upon Chloramphenicol Addition, Related to Figure 2**

(A) *ATP3* mRNA ratio of association to mitochondria was measured in WT, *sch9*Δ and *reg1*Δ mutant strains, and upon chloramphenicol addition (WT + Chlo.) (n>27). Error bar represents s.e.m.

(B-F) Mitochondrial volume fraction (B), cell volume (C), mitochondrial volume (D), adjusted mitochondrial volume fraction (E), and vacuole volume fraction (F) in WT, mutant, and chloramphenicol conditions (n>27).

(G) Predicted relationship of ratio of mRNA localization to mitochondria and non-adjusted mitochondrial volume fraction from mathematical modeling of WT strains was plotted as green dotted lines. Mean value of each axis of WT, mutants cells, and cells from chloramphenicol added conditions were plotted as cross. Error bar represents s.e.m.

(H, I) Relationship of the ratio of *ATP3* mRNA localization to mitochondria and mitochondrial volume fraction from mathematical modeling. (I) Mitochondrial volume fraction was adjusted with effective cytoplasm volume. Green lines represent the relationship in WT cells, orange, blue,and grey lines represent *sch9*Δ and *reg1*Δ mutant cells, and chloramphenicol added condition respectively. Lines were plotted as described in Methods.

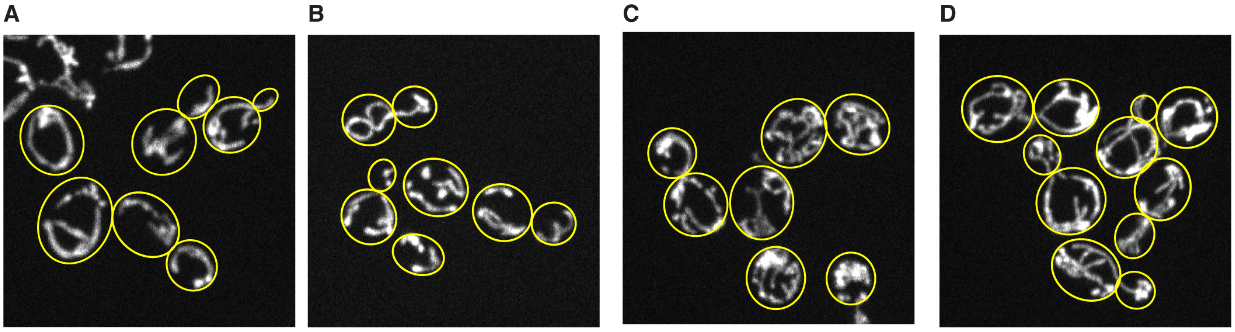


**Figure S6.** ***reg1*Δ Mutant Strains and Chloramphenicol Addition Increases Mitochondrial Volume Fraction, Related to Figure 2**

Z-projected image of mitochondria in live cells.

(A) Wild type cells grown in YPAD medium.

(B) *sch9*Δ mutant cells grown in YPAD medium.

(C) *reg1*Δ mutant cells grown in YPAD medium.

(D) Wild type cells grown in YPAD medium containing 1μg/ml of chloramphenicol.

Mitochondria were visualized by Su9-mCherry. Yellow lines represent cell boundary.

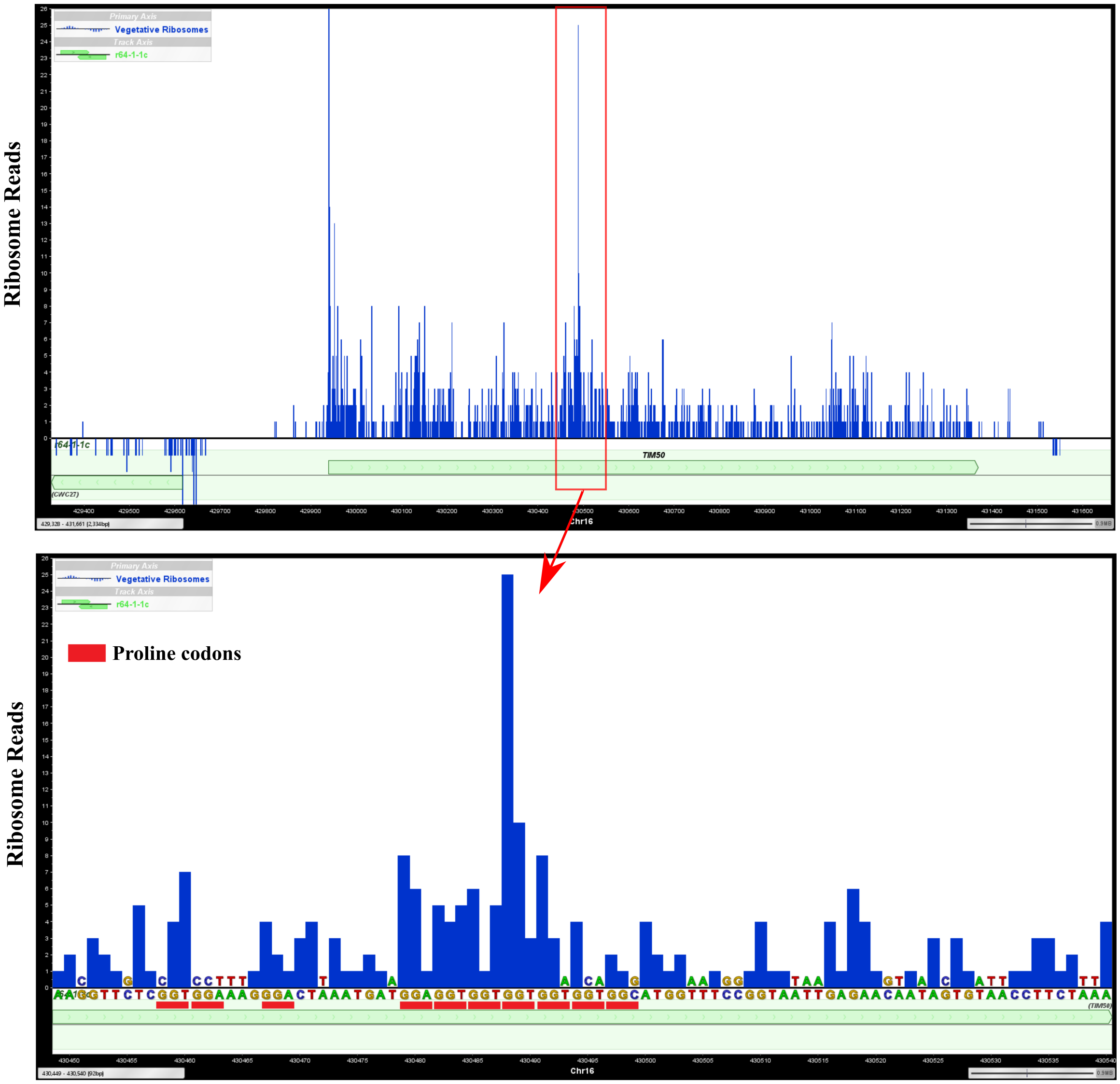


**Figure S7. Ribosomes are Enriched at a Polyproline Sequence of *TIM50* mRNA in a Previously Generated Ribosome Profiling Dataset, Related to Figure 5**

Proline codons were highlighted with red lines below *TIM50* DNA sequences. A site position of ribosome-covered fragments from a previous ribosome profiling dataset (Zid and O'Shea, 2014) are shown as ribosome reads.

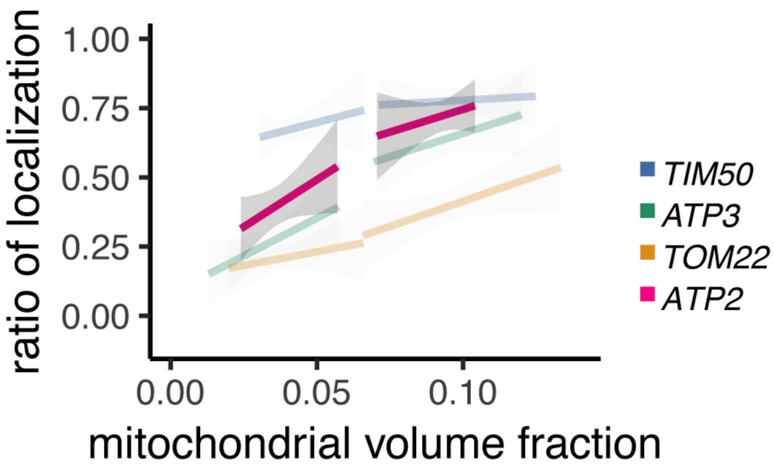


**Figure S8. Mitochondrial Volume Fraction Correlates with *ATP2* mRNA Localization, Related to Figure 5**

Relationship between the mitochondrial volume fraction and the ratio of *ATP2* mRNA (pink) localization to mitochondria with *TIM50*, *ATP3*, and *TOM22* mRNA, which are copied from Figure 1D. Trend line was depicted according to the best linear fit of the ratio of localization and mitochondrial volume fraction of single cells in each condition of different mRNAs (n>15). Gray regions surrounding the trend lines represent the 95 confidence interval (CI) for each line.

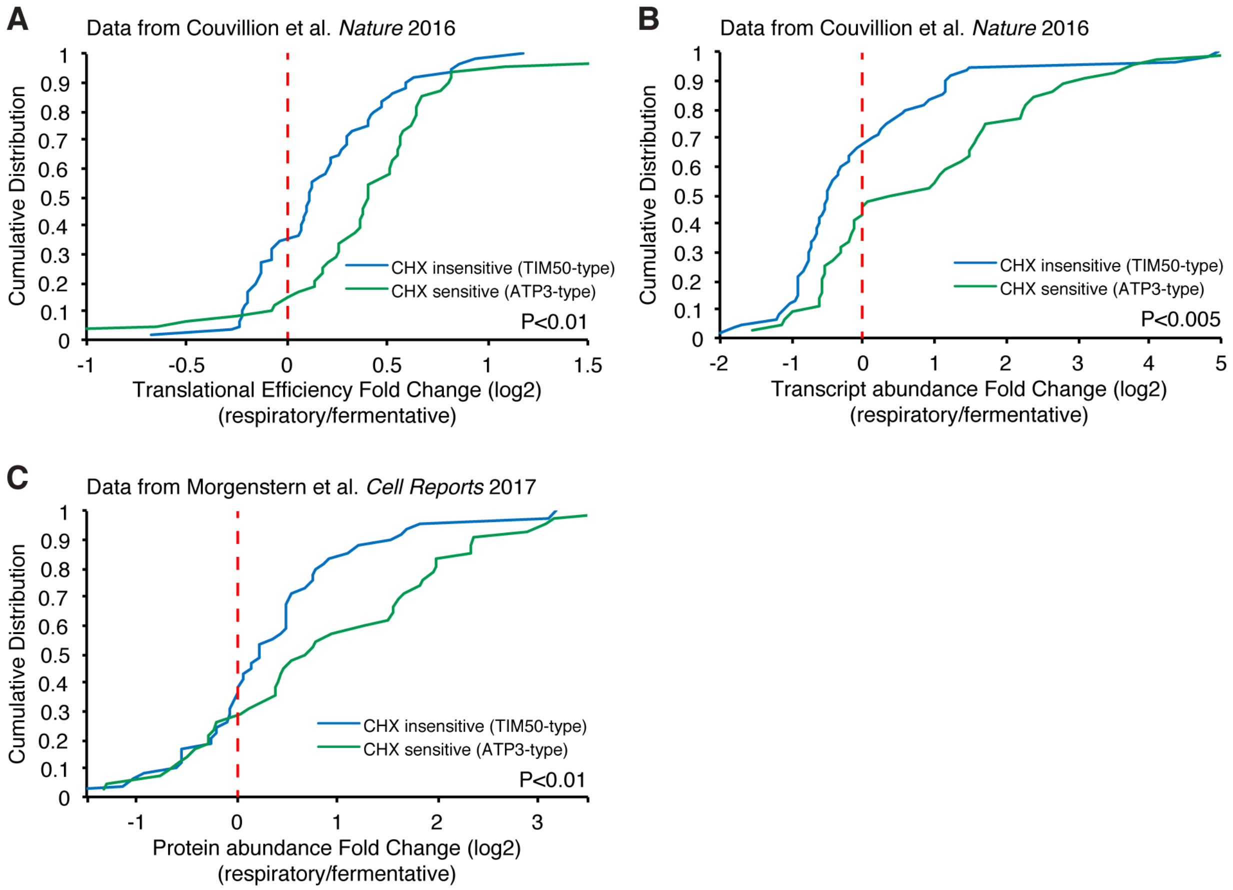


**Figure S9. Respiration Induced Increased Protein Production, Related to Figure 5**

(A, B) Translational efficiency (A) and transcript abundance (B) increase upon a switch from fermentative to respiratory conditions for ATP3-type genes. Cumulative distribution of translational efficiency and transcript abundance from previous ribosome profiling dataset (Couvillion et al., 2016) of TIM50-type and ATP3-type genes are depicted with blue and green lines, respectively. (C) Protein abundance increases upon switch from fermentative to respiratory conditions in ATP3-type genes. Cumulative distribution of protein abundance from previous mass spectrometry dataset (Morgenstern et al., 2017) of TIM50-type and ATP3-type genes are depicted with blue and green lines, respectively. Statistical significance in this figure was assessed by Student's t-test.

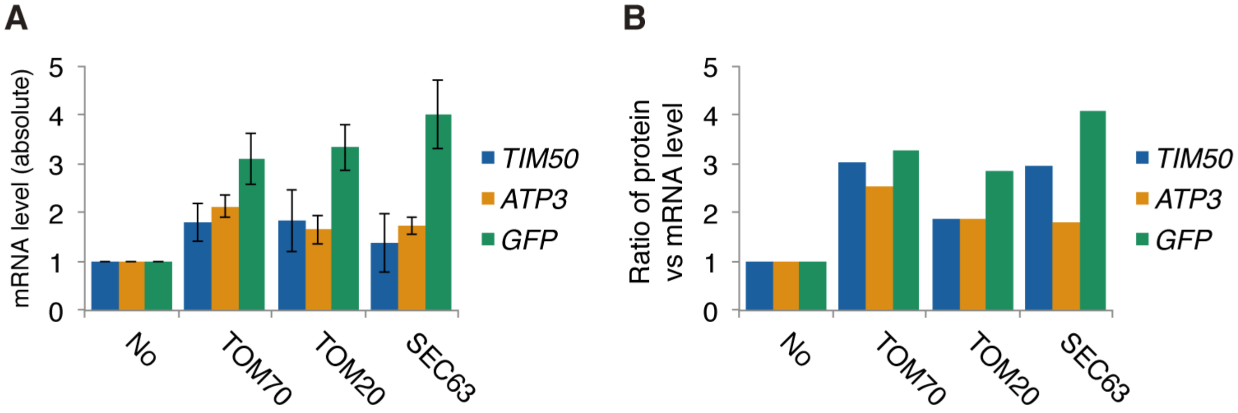


**Figure S10. Ratio of Protein Versus mRNA Expression Levels of Artificially Tethered Reporter mRNAs, Related to Figure 6**

(A) Expression level of reporter mRNAs tethered to mitochondria or ER. mRNA expression was analyzed using qPCR for *GFP*. Error indicates standard deviation of three independent experiments.

(B) Ratio of protein versus mRNA expression level of reporter mRNAs tethered to mitochondria (*TOM70* and *TOM20*) or ER (*SEC63*).

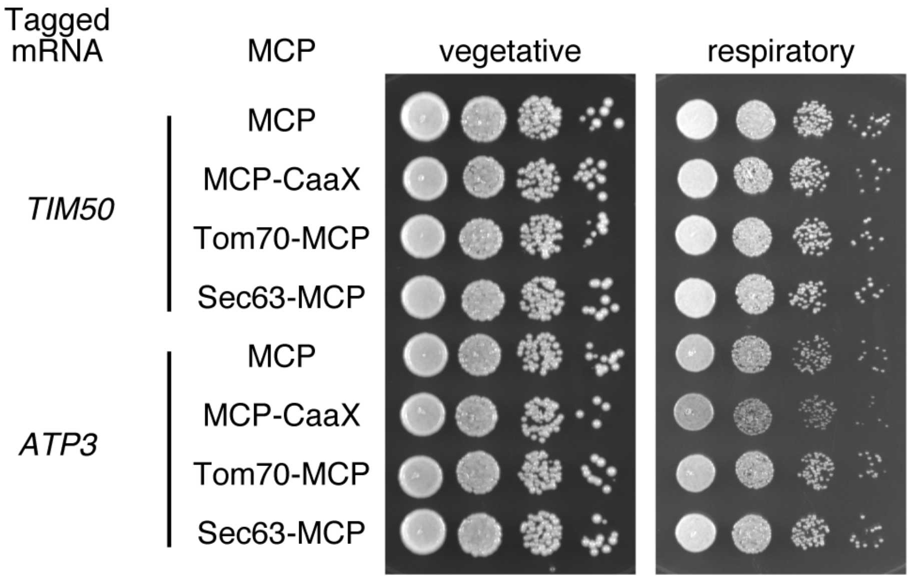


**Figure S11. Growth Assay for Strains with mRNA Tethered to Either Mitochondria or ER or mRNA Localized to the Plasma Membrane, Related to Figure 6**

*TIM50* and *ATP3* mRNAs were localized to the plasma membrane using CaaX-tag harboured MCP-GFP proteins, to mitochondria by tethering to TOM70-MCP, or to ER by tethering to Sec63-MCP. The growth was tested on YPAD (fermentative) and YPAGE (respiratory) conditions at 30°C for two days and three days, respectively. We took stationary phase cells and made 10x serial dilutions of them before plating.

### Table S1. Yeast strains and plasmids used in this study

| Strain/plasmids | Genotype[plasmid](plasmid number) | Source |
|---|---|---|
| Strains | | |
| W303-1a | MATa ade2-1 can1-100 his3-11 leu2-3 trp1-1 ura3 | Lab stock |
| TTY1160 | W303-1a TIM50-tag | This study |
| TTY584 | W303-1a ATP3-tag | This study |
| TTY439 | W303-1a TOM22-tag | This study |
| TTY1373 | W303-1a ATP1-tag | This study |
| TTY1374 | W303-1a ATP2-tag | This study |
| TTY1377 | W303-1a ATP7-tag | This study |
| Plasmids | | |
| pSH47 | Gal1p-Cre | Haim-Vilmovsky et al., 2009 |
| pLOXHIS5MS2L | | Haim-Vilmovsky et al., 2009 |
| pMS2CPGFP(x4) | | Haim-Vilmovsky et al., 2009 |
| pvt100-dsRed | | Lab stock |
| pFA6a-link-yomCherry-SpHis5 | | Lee et al., 2013 |
| pFA6a-link-yoGFP-SpHis5 | | Lee et al., 2013 |
| pFA6a-link-yoGFP- CaUra3 | | Lee et al., 2013 |
| pFA6a-hphMX6 | | Lab stock |
| NHB084 | IRFP-Kan | Gift from Nan Hao lab |
| TTP076 | pRS406*GPD*p-Su9-mCherry | This study |
| TTP080 | pRS405*CYC1*p-MS2-4xGFP | This study |
| TTP133 | pRS403*TIM50*-tag | This study |
| TTP134 | pRS403*ER-TIM50*-tag | This study |
| TTP135 | pRS403*TOM22*-tag | This study |
| TTP136 | pRS403*ER-TOM22*-tag | This study |
| TTP155 | pRS403*TIM50p-TIM50mts(1-300)-TIM50cds-flagyoGFP-TIM50ter-MS2tag* | This study |
| TTP161 | pRS403*TIM50p-ATP3mts(1-300)-ATP3cds-flagyoGFP-TIM50ter-MS2tag* | This study |
| TTP160 | pRS403*TIM50p-ATP3mts(1-300)-TIM50cds-flagyoGFP-TIM50ter-MS2tag* | This study |
| TTP162 | pRS403*TIM50p-TIM50mts(1-300)-ATP3cds-flagyoGFP-TIM50ter-MS2tag* | This study |
| TTP145 | pRS403*TIM50p-TIM50mts(1-300)-TIM50cds-flagiRFP-TIM50ter-MS2tag* | This study |
| TTP147 | pRS403*TIM50p-ATP3mts(1-300)-ATP3cds-flagiRFP-TIM50ter-MS2tag* | This study |
| TTP146 | pRS403*TIM50p-ATP3mts(1-300)-TIM50cds-flagiRFP-TIM50ter-MS2tag* | This study |
| TTP148 | pRS403*TIM50p-TIM50mts(1-300)-ATP3cds-flagiRFP-TIM50ter-MS2tag* | This study |
| TTP174 | pRS403*TIM50p-TIM50mts(1-300)-TIM50cds(delpolyP7)-flagyoGFP-TIM50ter-MS2tag* | This study |
| TTP179 | pRS403*TIM50p-TIM50mts(1-300)-TIM50cds(delpolyP7)-flagiRFP-TIM50ter-MS2tag* | This study |
| TTP158 | pRS403*TIM50p-flagyoGFP-TIM50ter-MS2tag* | This study |
| TTP119 | pFA6a-link-MCP-SpHis5 | This study |
| TTP153 | pFA6a-link-MCP-CaUra3 | This study |
| TTP167 | pRS405CYC1p-MS2-2xGFP-CaaX | This study |

### Table S2. List of oligonucleotides used for plasmid construction

| Gene Name | Oligonucleotides used for plasmid construction |
|---|---|
| *MS2-GFP(x4)* | 5'-GACTAGTCGGATGCAAGGGTTCGAATCCCTTAGCTCTC-3' |
| | 5'-CCGCTCGAGGGCCGCAAATTAAAGCCTTCGAGCGTCC-3' |
| *TIM50-Tag* | 5'-CCTTATTTGAAGAGGAAAAGAAAAAGAAGAAGATTGCTGAATCCAAATAAA ACGCTGCAGGTCGACAACCC-3' |
| | 5'-ACATACACACATAGATACGTAGATACATGAGAAGAGGGTTTACATGAAAAG CATAGGCCACTAGTGGATC-3' |
| *RPL2-Tag* | 5'-ATAGAATGAAAGTAAAAGACAGACCAAGAGGCAAAGATGCAAGATTATAAA ACGCTGCAGGTCGACAACCC-3' |
| | 5'-AACGAAGAAAAGGTCCCTATGTTGCTTCTTTTCTACTAAAATTAGCCTATGCA TAGGCCACTAGTGGATC-3' |
| *ATP3-Tag* | 5'-TTACTAATGAACTGGTTGATATTATTACTGGTGCTTCCTCTTTGGGATGAAAC GCTGCAGGTCGACAACCC-3' |
| | 5'-TTCTACAAAAACAACGTCAAATAAAGAGGCAATGCAGGGTGATTTTTTTAGC ATAGGCCACTAGTGGATC-3' |
| *TOM22-Tag* | 5'-ATAACATATTGGCCCAAGGTGAAAAAGATGCTGCAGCAACAGCCAATTAAA ACGCTGCAGGTCGACAACCC-3' |
| | 5'-ATGTATGGCTCCTTTTCTAAAACCCTCTCTTTTCTTTTACATCATTAAAAGCAT AGGCCACTAGTGGATC-3' |
| *ATP1-Tag* | *5'-GTTGGCATCTCTAAAGAGTGCTACTGAATCATTTGTTGCCACTTTTTAAAACGCTGCAGGTC GACAACCC-3'* |
| | *5'-ATTTCTTTTTGAGACGTACCTTATATTCATTTTTATTTTTTTAGTTCACAGCATAGGCCACTAG TGGATC-3'* |
| *ATP2-Tag* | *5'-TGAAGATGTTGTTGCTAAAGCTGAAAAGTTAGCCGCTGAAGCCAACTAGAACGCTGCAGGT CGACAACCC-3'* |
| | *5'-TACCTTCGGTATTTCAAATTTTGCTTCCCTTGGTTTAAGCTTTATTTCTTGCATAGGCCACTA GTGGATC-3'* |
| *ATP7-Tag* | *5'-GGACGTACCTGGTTACAAGGACAGATTCGGCAATTTGAATGTGATGTAGAACGCTGCAGGT CGACAACCC-3'* |
| | *5'-TGTGAAAAAAATAATAGAATATGGTGCGTAATATATAGAGGTAAAGGGTAGCATAGGCCACT AGTGGATC-3'* |
| *GPDp Sac2 F-680* | 5'-TCCCCGCGGCAGTTCGAGTTTATCATTATCAATACTGCC-3' |
| *GPDp Hind3 R-1* | 5'-CCCAAGCTTTTTGTTTGTTTATGTGTGTTTATTCGAAAC-3' |
| *color START Spe1 F* | 5'-GACTAGTCGGTGACGGTGCTGGTTTA-3' |
| *color ADHt Kpn1 R* | 5'-GGGGTACCTTACCCTGTTATCCCTAGCGGATCTGCCGG-3' |
| *C450_2C1_top* | 5'-ATGGATCCAGTTGTCGTATTGGGTTTATGTTTGTCTTGTTTGTTACTACTTTCT TTGTGGAAACAATCTTACGGTGGCGGAAAGTTA-3' |
| *C450_2C1_bot* | 5'-TAACTTTCCGCCACCGTAAGATTGTTTCCACAAAGAAAGTAGTAACAAACAA GACAAACATAAACCCAATACGACAACTGGATCCAT-3' |
| *TIM50 F-500 Spe1* | 5'-CGGTGGCGGCCGCTCTAGAACTAGTACTGTGTTGGCTTTAACTCTTTAAATTC TCC-3' |
| *TIM50 R++500 Xho1* | 5'-AATTGGGTACCGGGCCCCCCCTCGAGATGTGACGGCAGTTCCTGACCTGATA |

| | |
|---|---|
| | GTG-3' |
| *GA_TIM50_F129* | 5'-CAATCTTACGGTGGCGGAAAGTTACAAAAAGAAACAAAAGACGACAAGCCTAAATC-3' |
| *GA_TIM50_R-1* | 5'-ACCCAATACGACAACTGGATCCATTGCAAGCGGGTGATTTTTGGAAGTTTATTCTAGC-3' |
| *TOM22 F-500 Spe1* | 5'-CGGTGGCGGCCGCTCTAGAACTAGTCAAAAAGAGCTAATCAACTCCTTGAACTTAG-3' |
| *TOM22 R++500 Xho1* | 5'-AATTGGGTACCGGGCCCCCCCTCGAGGTTTACGTTTTAGATTACCAAAAAGGAAGCATAG-3' |
| *GA_TOM22_F1* | 5'-CAATCTTACGGTGGCGGAAAGTTAATGGTCGAATTAACTGAAATTAAAGACGATGTC-3' |
| *GA_TOM22_R-1* | *5'-ACCCAATACGACAACTGGATCCATTTGAATGATGCTTATTTTGGGGTATATAGTTCCG-3'* |
| *GA_TIM50_F562* | *5'-GAGCCACCTTTCCCTGATTTACTATACCAAAGGCCATTAACTCTTGTTATCACA-3'* |
| *TIM50_R540* | *5'-TAGTAAATCAGGGAAAGGTGGCTCTTGGA-3'* |
| *yeGFP-CaaX_F* | *5'-GGTAAGGCTAGCGGTAAAAAGAAGAAAAAGAAGTCAAAGACAAAGTGTGTAATTATGTAACTGGTCGAGTCATGTAATTAGTTATGTC-3'* |
| *yeGFP-CaaX_R* | *5'-CTTTTTACCGCTAGCCTTACCAGCACTGCCTGCGCTATCGCTACCTTTGTACAATTCATCCATACCATGGGT-3'* |

**Table S3. mRNAs of conserved ATP synthetase components are localized to mitochondria in respiratory and CHX conditions.**

<table>
<tr><th rowspan="2"></th><th rowspan="2">Complex</th><th rowspan="2">Subunit</th><th rowspan="2">Gene</th><th colspan="2">mitochondrial localization</th><th rowspan="2">respiratory specific localization</th></tr>
<tr><th>-CHX</th><th>+CHX</th></tr>
<tr><td rowspan="6">conserved from bacteria to eukaryote</td><td rowspan="5">F1</td><td>α</td><td>ATP1</td><td>no</td><td>yes</td><td>yes</td></tr>
<tr><td>β</td><td>ATP2</td><td>no</td><td>yes</td><td>yes</td></tr>
<tr><td>γ</td><td>ATP3</td><td>no</td><td>yes</td><td>yes</td></tr>
<tr><td>OSCP/δ</td><td>ATP5</td><td>no</td><td>yes</td><td>NA</td></tr>
<tr><td>ε</td><td>ATP16</td><td>no</td><td>no</td><td>NA</td></tr>
<tr><td>F0</td><td>b</td><td>ATP4</td><td>no</td><td>yes</td><td>NA</td></tr>
<tr><td>non-conserved</td><td colspan="3">all the others</td><td>no</td><td>no</td><td>NA(ATP7: no)</td></tr>
<tr><td>Rühle et al., 2015</td><td colspan="3"></td><td colspan="2">Williams et al., 2014</td><td>This study</td></tr>
</table>

ATP complex gene conservation was obtained from Rühle and Leister, 2015 and we obtained data of CHX-dependent mitochondrial localization from proximity ribosomal profiling data (Williams et al., 2014). The CHX-dependent localization data was consistent with our study. Respiratory-specific localization was determined in this study.

### References

Arganda-Carreras, I., Kaynig, V., Rueden, C., Eliceiri, K.W., Schindelin, J., Cardona, A., and Sebastian Seung, H. (2017). Trainable Weka Segmentation: a machine learning tool for microscopy pixel classification. Bioinformatics *33*, 2424–2426.

Bertrand, E., Chartrand, P., Schaefer, M., Shenoy, S.M., Singer, R.H., and Long, R.M. (1998). Localization of ASH1 mRNA Particles in Living Yeast. Mol. Cell *2*, 437–445.

Chan, Y.H.M., Reyes, L., Sohail, S.M., Tran, N.K., and Marshall, W.F. (2016). Organelle Size Scaling of the Budding Yeast Vacuole by Relative Growth and Inheritance. Curr. Biol. *26*, 1221–1228.

Couvillion, M.T., Soto, I.C., Shipkovenska, G., and Churchman, L.S. (2016). Synchronized mitochondrial and cytosolic translation programs. Nature *533*, 499–503.

Firth, D. (1992). Bias reduction, the Jeffreys prior and GLIM. In Advances in GLIM and Statistical Modelling, L. Fahrmeir, B. Francis, R. Gilchrist, and G. Tutz, eds. (New York, NY: Springer New York), pp. 91–100.

Firth, D. (1993). Bias Reduction of Maximum Likelihood. Biometrika.

Gadir, N., Haim-Vilmovsky, L., Kraut-Cohen, J., and Gerst, J.E. (2011). Localization of mRNAs coding for mitochondrial proteins in the yeast Saccharomyces cerevisiae. RNA *17*, 1551–1565.

Haim-Vilmovsky, L., and Gerst, J.E. (2009). m-TAG: a PCR-based genomic integration method to visualize the localization of specific endogenous mRNAs in vivo in yeast. Nat. Protoc. *4*,

1274–1284.

Hocine, S., Raymond, P., Zenklusen, D., Chao, J.A., and Singer, R.H. (2012). Single-molecule analysis of gene expression using two-color RNA labeling in live yeast. Nat. Methods *10*, 119–121.

Jorgensen, P., Edgington, N.P., Schneider, B.L., Rupes, I., Tyers, M., and Futcher, B. (2007). The Size of the Nucleus Increases as Yeast Cells Grow. Mol. Biol. Cell.

Lee, S., Lim, W.A., and Thorn, K.S. (2013). Improved Blue, Green, and Red Fluorescent Protein Tagging Vectors for S. cerevisiae. PLoS One *8*, 4–11.

Morgenstern, M., Stiller, S.B., Lübbert, P., Peikert, C.D., Dannenmaier, S., Drepper, F., Weill, U., Höß, P., Feuerstein, R., Gebert, M., et al. (2017). Definition of a High-Confidence Mitochondrial Proteome at Quantitative Scale. Cell Rep. *19*, 2836–2852.

Rafelski, S.M., Viana, M.P., Zhang, Y., Chan, Y.-H.M., Thorn, K.S., Yam, P., Fung, J.C., Li, H., Costa, L.D.F., and Marshall, W.F. (2012). Mitochondrial network size scaling in budding yeast. Science *338*, 822–824.

Rühle, T., and Leister, D. (2015). Assembly of F1F0-ATP synthases. Biochim. Biophys. Acta *1847*, 849–860.

Shah, P., Ding, Y., Niemczyk, M., Kudla, G., and Plotkin, J.B. (2013). Rate-limiting steps in yeast protein translation. Cell *153*, 1589–1601.

Tinevez, J.Y., Perry, N., Schindelin, J., Hoopes, G.M., Reynolds, G.D., Laplantine, E., Bednarek,

S.Y., Shorte, S.L., and Eliceiri, K.W. (2017). TrackMate: An open and extensible platform for single-particle tracking. Methods.

Tsuboi, T., Kuroha, K., Kudo, K., Makino, S., Inoue, E., Kashima, I., and Inada, T. (2012). Dom34: Hbs1 Plays a General Role in Quality-Control Systems by Dissociation of a Stalled Ribosome at the 3' End of Aberrant mRNA. Mol. Cell *46*, 518–529.

Tutucci, E., Vera, M., Biswas, J., Garcia, J., Parker, R., and Singer, R.H. (2018a). An improved MS2 system for accurate reporting of the mRNA life cycle. Nat. Methods *15*, 81–89.

Tutucci, E., Vera, M., and Singer, R.H. (2018b). Single-mRNA detection in living S. cerevisiae using a re-engineered MS2 system. Nat. Protoc. *13*, 2268–2296.

Viana, M.P., Lim, S., and Rafelski, S.M. (2015). Quantifying mitochondrial content in living cells (Elsevier Ltd).

Williams, C.C., Jan, C.H., and Weissman, J.S. (2014). Targeting and plasticity of mitochondrial proteins revealed by proximity-specific ribosome profiling. Science (80-. ). *346*, 748–751.

Zid, B.M., and O'Shea, E.K. (2014). Promoter sequences direct cytoplasmic localization and translation of mRNAs during starvation in yeast. Nature *514*, 117–121.
